# Characterization of small molecule induced changes in Parkinson’s-related trafficking via the Nedd4 ubiquitin signaling cascade

**DOI:** 10.1101/2020.06.01.128348

**Authors:** A. Katherine Hatstat, Hannah D. Ahrendt, Matthew W. Foster, Leland Mayne, M. Arthur Moseley, S. Walter Englander, Dewey G. McCafferty

## Abstract

The benzdiimidazole NAB2 rescues α-synuclein-associated trafficking defects associated with early onset Parkinson’s disease in a Nedd4-dependent manner. Despite identification of E3 ubiquitin ligase Nedd4 as a putative target of NAB2, its molecular mechanism of action has not been elucidated. As such, the effect of NAB2 on Nedd4 activity and specificity was interrogated through biochemical, biophysical, and proteomic analyses. NAB2 was found to bind Nedd4 (K_D_^*app*^ = 42 nM), but this binding is side chain mediated and does not alter its conformation or ubiquitination kinetics *in vitro*. Nedd4 co-localizes with trafficking organelles, and NAB2 exposure did not alter its colocalization. Ubiquitin-enrichment coupled proteomics revealed that NAB2 stimulates ubiquitination of trafficking and transport associated proteins, most likely through modulating the substrate specificity of Nedd4, providing a putative protein network involved in the NAB2 mechanism.

## Introduction

Parkinson’s disease (PD), a neurodegenerative disorder characterized by distinct motor and non-motor symptoms, results from targeted loss of dopaminergic neurons in the midbrain. Neurodegeneration can result from many causes including cellular toxicity induced by aggregation of toxic proteins. In one type of PD, mutations, duplications or triplications at the *SNCA* gene locus induce production of toxic proteoforms of α-synuclein.^1–4^ In its wild-type form, α-synuclein is a membrane associated protein involved in trafficking and transport processes. In the presence of PD-associated *SNCA* alterations, however, α-synuclein forms toxic aggregates that induce generation of reactive oxygen species and disrupt cellular processes including trafficking and transport.^5–8^

There is increased interest in developing treatments for PD that target disrupted cellular processes associated with neuronal toxicity and neurodegeneration to support front line dopamine replacement therapy. Pharmacological alleviation of neurotoxicity could prevent further neurodegeneration and help mitigate some disease related sequelae such as reduced sensitivity to levodopa over extended periods of use.^9–12^ Identifying critical neurotoxicity signaling players has led to the development of cellular models to enable phenotype-driven screening for identification of targets and potential small molecule leads to reverse the cellular toxicity signaling associated with neurodegeneration. Recently, the Lindquist group developed a yeast-based phenotypic model of α-synuclein toxicity, and discovered a small molecule *N*-arylbenzdiimidazole (NAB) that significantly alleviated most of the major phenotypic markers of α-synuclein toxicity (Figure 1A).^1,13^ Counter genetic screening revealed that the activity of NAB analogs (NABs) was dependent the yeast protein Rsp5, an E3 ubiquitin ligase. Further validation in mammalian cell models indicated that NAB activity was conserved and was shown to be dependent upon Nedd4, the mammalian homolog of Rsp5. Finally, phenotype-driven structure activity relationship (SAR) optimization of the NAB scaffold afforded a derivative, NAB2, with improved activity over the lead compound (Figure 1A).^1,13^ However, since these initial studies, the molecular mechanism of action of NAB2 and the role of Nedd4 in the alleviation of toxicity have yet to further elucidated.

**Figure 1.**
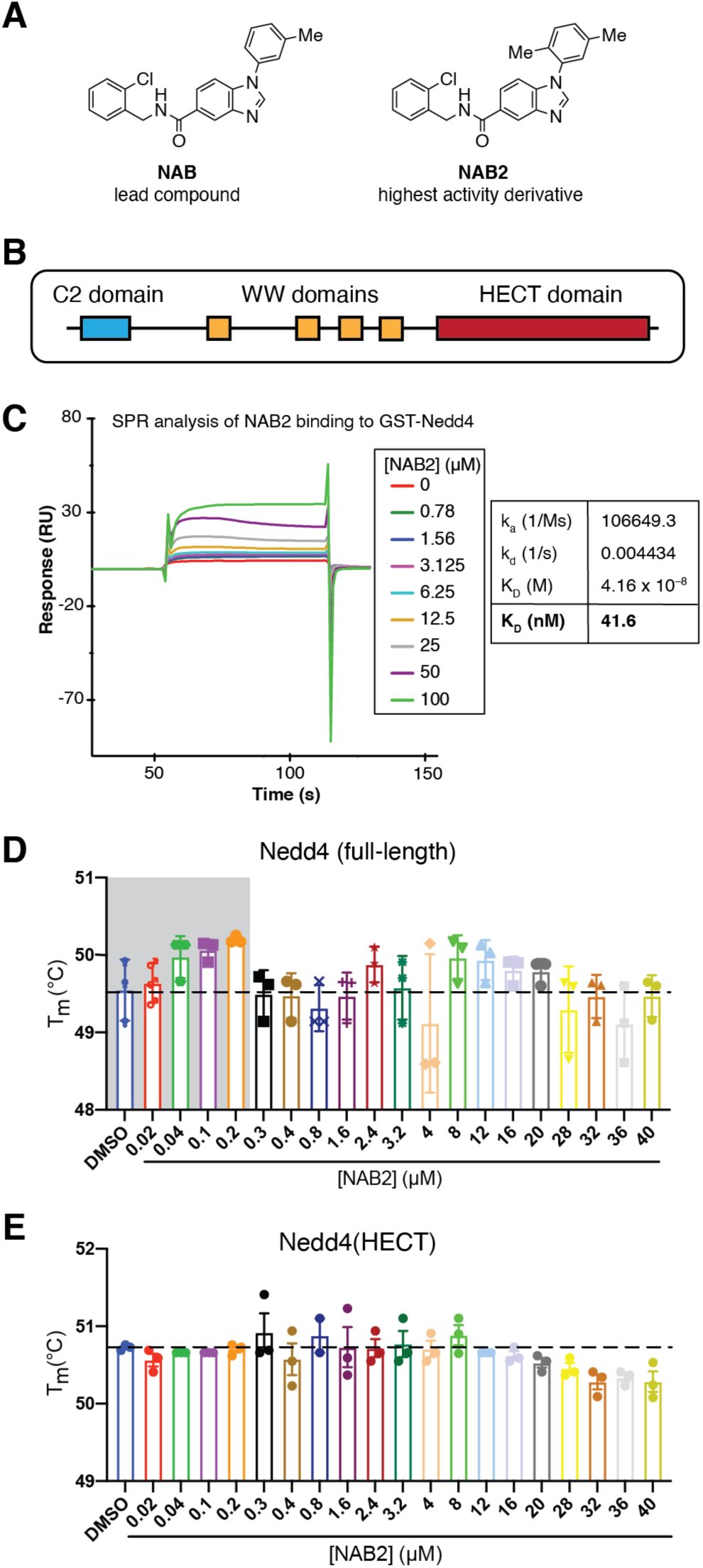
*In vitro* analyses demonstrate NAB2 engagement with Nedd4. **(A)** Phenotypic screening identified a lead compound of an *N-* arylbenzdiimidazole scaffold (NAB) which significantly alleviated markers of α-synuclein toxicity (EC_40_ = 7.5 *µ*M). Further optimization of the lead compound provided access to NAB2 as the most active derivative with an EC_40_ = 4.5 *µ*M in phenotype-driven assays.^2^ **(B)** Linear representation of full-length Nedd4 depicts the subdomains of Nedd4. The enzyme contains a C2 domain (cyan) which signals for membrane localization, four WW domains (yellow) for substrate recognition, and a catalytic HECT domain (red). For *in vitro* analyses, full-length Nedd4 and the isolated Nedd4(HECT) domain were used. **(C)** Characterization of NAB2 binding by surface plasmon resonance (SPR) for low molecular weight kinetics show that NAB2 binds to GST-Nedd4 with an apparent affinity of 41.6 nM relative to a control surface. **(D**,**E)** To complement SPR with a thermodynamic measurement of ligand binding, protein thermal shift assays (PTSA) were conducted using SYRPO orange detection of Nedd4 stability over a range of NAB2 concentrations. Nedd4 (4 *µ*M) was incubated with NAB2 (concentration indicated on graph) for 30 minutes. SYPRO orange was added and samples were analyzed using a Roche LightCycler 480 qPCR instrument. Samples were heated from 20 to 85 °C, and T_m_ was determined as the absolute minimum of the negative first derivative of the melting curve. PTSA analysis shows **(D)** a NAB2-dependent change in full-length Nedd4 stability at nanomolar concentrations (highlight in grey) but is not reflective of a single binding event, whereas **(E)** Nedd4(HECT) shows little change until a slight de-stabilization at higher micromolar concentrations.

The identification of Nedd4 as a potential target in PD-associated toxicity is intriguing for a number of reasons. First, Nedd4 has previously been implicated in cellular responses to PD-associated toxicity as it was shown to interact with and ubiquitinate α-synuclein, resulting in ubiquitination-induced degradation of the protein.^14^ Second, Nedd4 ubiquitinates aggregated α-synuclein more extensively than monomeric α-synuclein, indicating that it has potential to be specifically involved in clearance of the toxic proteoform.^15^ Additionally, Nedd4 has also been recently implicated in cellular responses to stressors including heat shock, oxidative stress, and protein misfolding.^16–18^ As α-synuclein-associated toxicity is a result of both protein aggregation and induced oxidative stress, the implication of Nedd4 in the response to α-synuclein-specific toxicity is promising.

Despite the potential role of Nedd4 in cellular responses to α-synuclein toxicity, the enzyme is considered a non-canonical drug target due to its lack of discrete active site and fairly complex enzymatic mechanism of activation, substrate recognition, and ubiquitination transfer. As a HECT-type E3 ubiquitin ligase (Figure 1B), Nedd4 depends upon protein-protein interactions with the upstream E2 conjugating enzyme and downstream substrate. It requires two chemical steps: transthioesterification for receipt of ubiquitin from the E2 enzyme, and isopeptide bond formation for passage of ubiquitin to the substrate. Despite its complex activation mechanism, as an E3 ligase Nedd4 is believed to be the only member of this signaling cascade that directly interacts with substrates, and thus confers the greatest specificity for chemical manipulation of ubiquitination. While there is some precedence for small molecule inhibition or alteration of Nedd4 activity or processivity,^19–21^ ligands identified to date that specifically target the ligase are primarily covalent modifiers, and the ability to target Nedd4 with non-covalent ligands remains underexplored. We therefore sought to interrogate the NAB mechanism to better understand its impact on the activity and specificity of Nedd4, specifically as it pertains to α-synuclein toxicity.

Preliminary characterizations of the mechanism of NABs indicate that, while activity is dependent upon Rsp5/Nedd4, treatment with NAB alleviates toxicity independent of α-synuclein levels.^1,13^ Additionally, NAB treatment did not inhibit Rsp5/Nedd4-dependent ubiquitination *in vitro*. Despite the lack of inhibition, a single point mutation in Rsp5 abolished NAB activity.^1^ We hypothesize that NAB may modulate the Nedd4 activity (i.e. kinetics or ubiquitin linkage specificity), or may alter its substrate specificity or interactome, thereby influencing Nedd4-dependent target ubiquitination and in turn impacting downstream toxicity signaling events. To interrogate these hypotheses, we employed *in vitro* biochemical and biophysical analyses in combination with cellular and proteomic experiments to characterize the mechanism of Nedd4 in response to treatment with NAB2, the highest activity NAB derivative.^1^ Herein, we present the results of these analyses which indicate that NAB2 engages with Nedd4 with apparent affinity in the nanomolar range but does not induce changes in Nedd4 activity, conformation or ubiquitin linkage specificity *in vitro*. At a cellular level, we observed small but significant NAB2-dependent changes in Nedd4 co-localization with trafficking organelles. Through proteomic analyses, we demonstrate that α-synuclein toxicity drastically remodels the ubiquitylome. Further, we identify NAB2-dependent changes in protein ubiquitination. Specifically, NAB2 treatment induces significant changes in ubiquitination of trafficking and transport associated proteins relative to control samples. Together, these experiments demonstrate that NAB2 engages with Nedd4 and identify NAB2-dependent changes in ER to Golgi transport, providing insight into the NAB2 mechanism and a putative protein network affected by NAB2 treatment.

## Results

### Surface plasmon resonance and protein thermal shift assays to characterize NAB2 binding to Nedd4

Prior to *in vitro* analyses of NAB2 mechanism of action or Nedd4 enzymology, we first sought access to the ligand for use as a chemical probe and to Nedd4 and associated enzymes as recombinant proteins. Specifically, we synthetically accessed NAB2 according to the synthetic route previously described (Figure 1A; Supplemental Scheme 1).^1^ Additionally, protein expression and purification strategies were established to allow facile access to Nedd4 in various forms (full-length enzyme and isolated catalytic HECT domain) and to Ubch5a, an E2 conjugating enzyme that interacts with Nedd4.^22^ Enzymes were accessed as recombinant proteins by expression in *E. coli* and were subsequently purified through optimized affinity purification strategies (described in Methods). The activity of the purified enzymes was confirmed by immunoblotting-based detection of Nedd4 autoubiquitination or ubiquitin chain formation (Supplemental Figure 1). Access to the ubiquitin signaling cascade as recombinant proteins enabled subsequent *in vitro* analyses.

Initial characterization of NAB2 revealed that its activity was dependent upon Rsp5 or homolog Nedd4 in yeast and mammalian models, respectively.^1,13^ Therefore, we first sought to characterize the on-target binding of NAB2 through *in vitro* analyses of ligand engagement. To quantify the affinity of NAB2 to Nedd4, ligand binding was characterized through surface plasmon resonance (SPR) using GST-Nedd4. Following protein immobilization and equilibration, the chip surface was exposed to a concentration gradient of NAB2 from 0 *µ*M to 100 *µ*M for characterization of low molecular weight kinetics. SPR analysis of binding relative to a control surface indicated that NAB2 bound to GST-Nedd4 with an apparent K_D_ (K_D_^*app*^) of 41.6 nM (Figure 1C). Further analysis by SPR of ligand binding to the isolated catalytic HECT domain, a stable subdomain of the full-length ligase, did not show a canonical binding curve (Supplemental Figure 2a). This result indicates that NAB2 binding is presumably occurring upstream of the C-terminus catalytic HECT domain. To further validate binding analyses and to investigate the stoichiometry of the NAB2/Nedd4 interaction, we turned to thermodynamic measurements of ligand binding. Using isothermal titration calorimetry (ITC), no binding event was detected upon titration of NAB2 into Nedd4(HECT) domain (Supplemental Figure 2b), validating the results obtained by SPR. ITC analysis of full-length Nedd4 was unsuccessful as the full-length protein suffered from instability and precipitation during extended ITC titrations (Supplemental Figure 2c), so we instead employed a protein thermal shift assay (PTSA) to further characterize ligand binding to Nedd4 (Figure 1D). PTSA analyses reveal a NAB2-dependent shift in the T_m_ of full-length Nedd4 at nanomolar concentrations, indicating a change in protein stability. While this result is consistent with the K_D_^*app*^ determined by SPR, the thermal shift results at higher NAB2 concentrations are not consistent as there is not a direct concentration-dependent thermodynamic shift across the full range of NAB2 concentrations tested. Instead, the stabilizing effect seen at low concentrations fluctuates, indicating that a more complex binding event may be occurring. PTSA analysis of NAB2-induced changes in Nedd4(HECT) thermostability are consistent with SPR and ITC results (Figure 1E). Cumulatively, *in vitro* analyses of target engagement demonstrate that NAB2 binds to Nedd4 with high apparent affinity upstream of the HECT domain. To further elucidate the mode of action of NAB2, we turned to *in vitro* biophysical and biochemical characterizations to identify NAB2-dependent alterations of Nedd4 conformation and activity.

### Bottom-up hydrogen-deuterium exchange mass spectrometry enables characterization of ligand-induced conformational changes

As SPR and PTSA results are indicative of a NAB2/Nedd4 binding event, we sought to characterize the ligand binding site and any NAB2-dependent conformational changes induced by NAB2 binding to Nedd4. This is particularly important in the case of Nedd4 as its enzymatic activity is governed by regulatory intramolecular interactions; thus, ligand-induced conformational changes may have functional implications.^23–27^ Due to the large size (∼104 kDa) and flexibility of full-length Nedd4 (Figure 2A), we were unable to pursue techniques like saturation transfer difference nuclear magnetic resonance (STD-NMR) or structural analyses like x-ray crystallography or cryo-EM to characterize the ligand/protein complex. We turned, instead, to bottom-up hydrogen-deuterium exchange mass spectrometry (HDX-MS).^28^ In this case, HDX-MS is useful for this application as it enables characterization of the relative order of a protein based on the rate of labile backbone proton exchange in the presence of deuterium, and the large size of Nedd4 is not a limitation. Further, there is precedence for the use of HDX-MS for characterization of protein-ligand complexes, and it has particular utility for ligands that induce conformational changes (i.e. calmodulin and Ca^2+^).^29–31^

**Figure 2.**
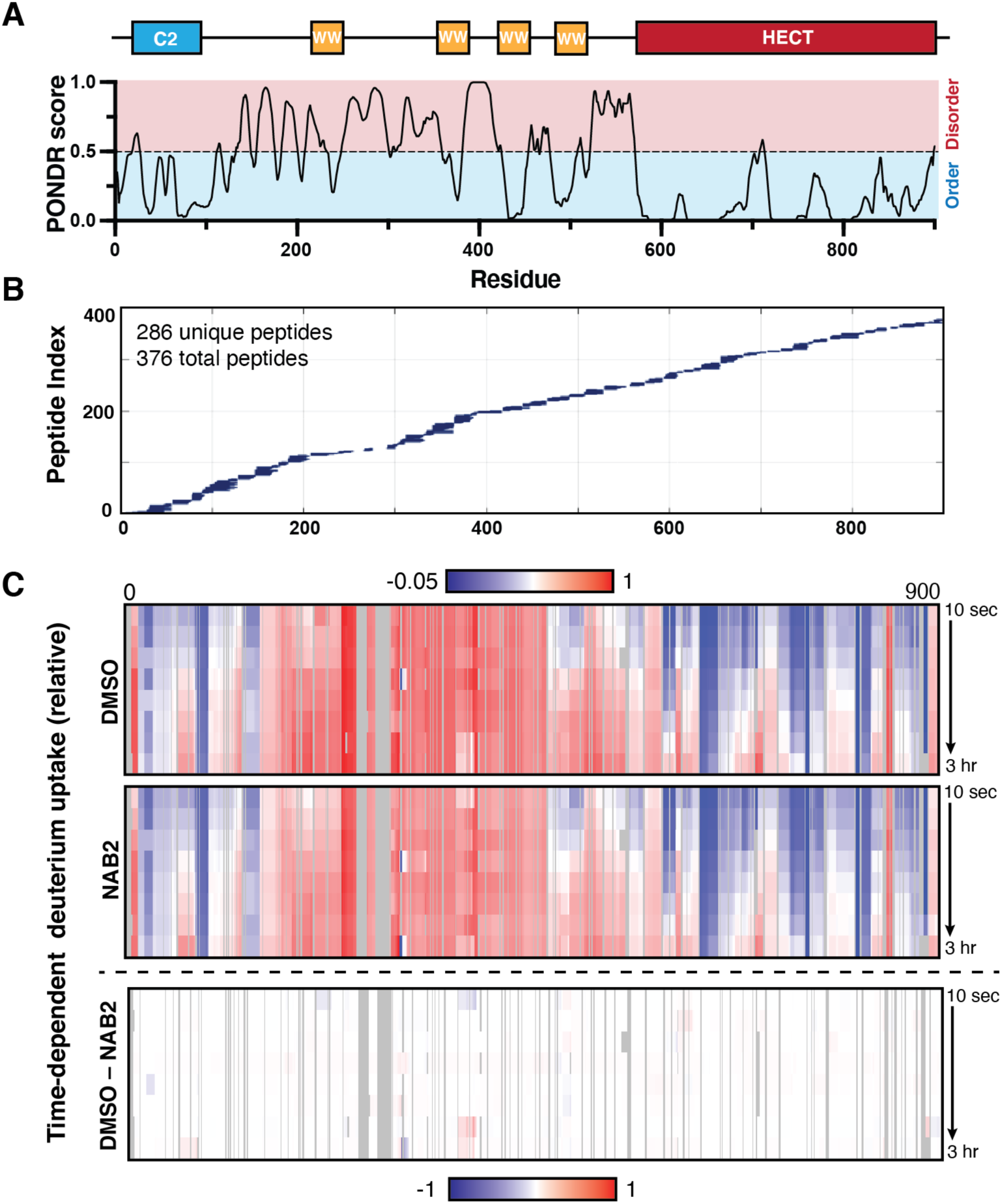
Hydrogen-deuterium exchanged coupled-mass spectrometry (HDX-MS) shows no significant ligand-induced changes Nedd4 conformation. As a multi-domain protein, Nedd4 contains regions of order and highly disordered linker regions. **(A)** The relative order of the protein and its subdomains was predicted with the PONDR algorithm (www.pondr.com) and corresponds well with the defined sub-domains of the ligase structure (top). **(B)** Peptide coverage map of full-length Nedd4 from diagnostic pepsin digest, which yielded 286 unique peptides and gave high coverage of the sequence for HDX analysis. **(C)** Nedd4 conformation was mapped by time-resolved HDX-MS wherein full-length Nedd4 (50 *µ*M) was incubated with NAB2 (500 *µ*M) or DMSO control. Samples were diluted 10-fold into deuterated buffer and incubated for the designated time prior to quenching to pH 2.4. Quenched samples were subjected to sequential, low-temperature (4 °C) pepsin digest, desalting, and LC-MS/MS. The degree of deuterium uptake in each peptide was compared to an “all-hydrogen” control and deuterium uptake across the Nedd4 sequence was mapped as a function of time using ExMS2. The rate of deuterium uptake into the Nedd4 backbone over time was consistent with the predicted order of the protein (top, compared to **A**). NAB2 treatment (middle) did not induce a significant conformational change relative to a DMSO-treated control as indicated by little difference in the subtraction map (bottom).

We hypothesized that if NAB2 binding shields the Nedd4 backbone we would detect a localized change in deuterium uptake. Further, if NAB2 treatment alters Nedd4 conformation by altering relative order of the enzyme or changing intramolecular interactions between Nedd4 subunits, the ligand-induced change could be detected by a change in the rate of deuterium uptake relative to a DMSO-treated control. To this end, HDX-MS analysis was conducted using recombinant, full-length Nedd4 in the presence of NAB2 or DMSO control. Prior to time-course analysis, peptide coverage of the protein sequence was confirmed via diagnostic pepsin digestion, LC-MS/MS, and automated database searching using SEQUEST. Test digestions revealed that pepsin digestion provided 376 total peptides (with 286 unique peptides), giving high coverage across the length of Nedd4 (Figure 2B). Control samples for “all hydrogen” and “all deuterium” conditions were additionally prepared by overnight incubation of Nedd4 with normal or deuterated buffer, respectively. Finally, a HDX time-course experiment was conducted at timepoints ranging from 10 seconds to 3 hours to enable exchange to occur in both disordered and ordered regions of the protein (Figure 2C). Over the time-course experiments, HDX data was collected at every timepoint for 752 of 900 residues, covering 83.5% of the total Nedd4 sequence. The results of bottom-up HDX-MS indicate that Nedd4 conformation is consistent with the relative order of the protein as predicted by the Predictor of Naturally Disordered Regions (PONDR) algorithm (Figure 2A; www.pondr.com) wherein more ordered regions of Nedd4 (N-terminal C2 domain and C-terminal HECT domain) did not uptake deuterium as rapidly as disordered linker regions. While the rate of deuterium uptake corresponds well with the predicted order of the Nedd4 structure (Figure 2C, top), the results show that NAB2 treatment (Figure 2C, middle) did not induce a significant global difference in the rate of deuterium uptake relative to the DMSO control (Figure 2C, bottom). This result indicates that NAB2 treatment does not induce a global change in Nedd4 conformation. It is important to note that a significant portion of the protein (between the C2 and HECT domain, encompassing the WW domains and linker regions) was partially or fully deuterated even at the shortest timepoint of the HDX-MS time-course experiment. If NAB2 induces a conformational change in this region, it would not be easily detected due to limitations in the sensitivity of the method employed unless the change resulted in significant shielding. We did, however, observe slight alterations in deuterium uptake in the C2, linker, and WW domain 2. This result is consistent with NAB2 binding upstream of the C-terminal HECT domain (as shown via SPR and PTSA), but further investigation of the NAB2 binding mode is needed and will be explored in future analyses. Cumulatively, the results from HDX-MS with biophysical and thermodynamic measurements of NAB2 binding show that NAB2 binding does not alter Nedd4 conformation *in vitro*. Nonetheless, we next sought to determine if NAB2 binding alters Nedd4 kinetics or ubiquitin linkage specificity.

### Quantitative MALDI-TOF and immunoblotting assays for measurement of NAB2-induced changes in Nedd4 activity

The initial characterizations of NAB2 by *in vitro* immunoblotting assays suggested that it did not inhibit Nedd4 activity.^1^ It was not determined, however, if NAB2 treatment altered Nedd4 kinetics or ubiquitin linkage specificity. Therefore, we sought to characterize NAB2-dependent changes of Nedd4 enzymology more thoroughly. To assay Nedd4 ubiquitin ligase activity in a sensitive and quantitative manner, we adapted a MALDI-TOF ubiquitination assay that measures monoubiquitin consumption over time relative to an internal standard.^32^ Compared to immunoblotting-based assays, this platform enables high-throughput, time-resolved screening of Nedd4 activity (Figure 3A,B). Analysis of relative Nedd4 activity by MALDI-TOF showed that NAB2 treatment did not significantly alter the rate of monoubiquitin consumption by the full-length ligase *in vitro* (Figure 3C), suggesting that NAB2 does not alter the kinetics of Nedd4-dependent ubiquitination.

**Figure 3.**
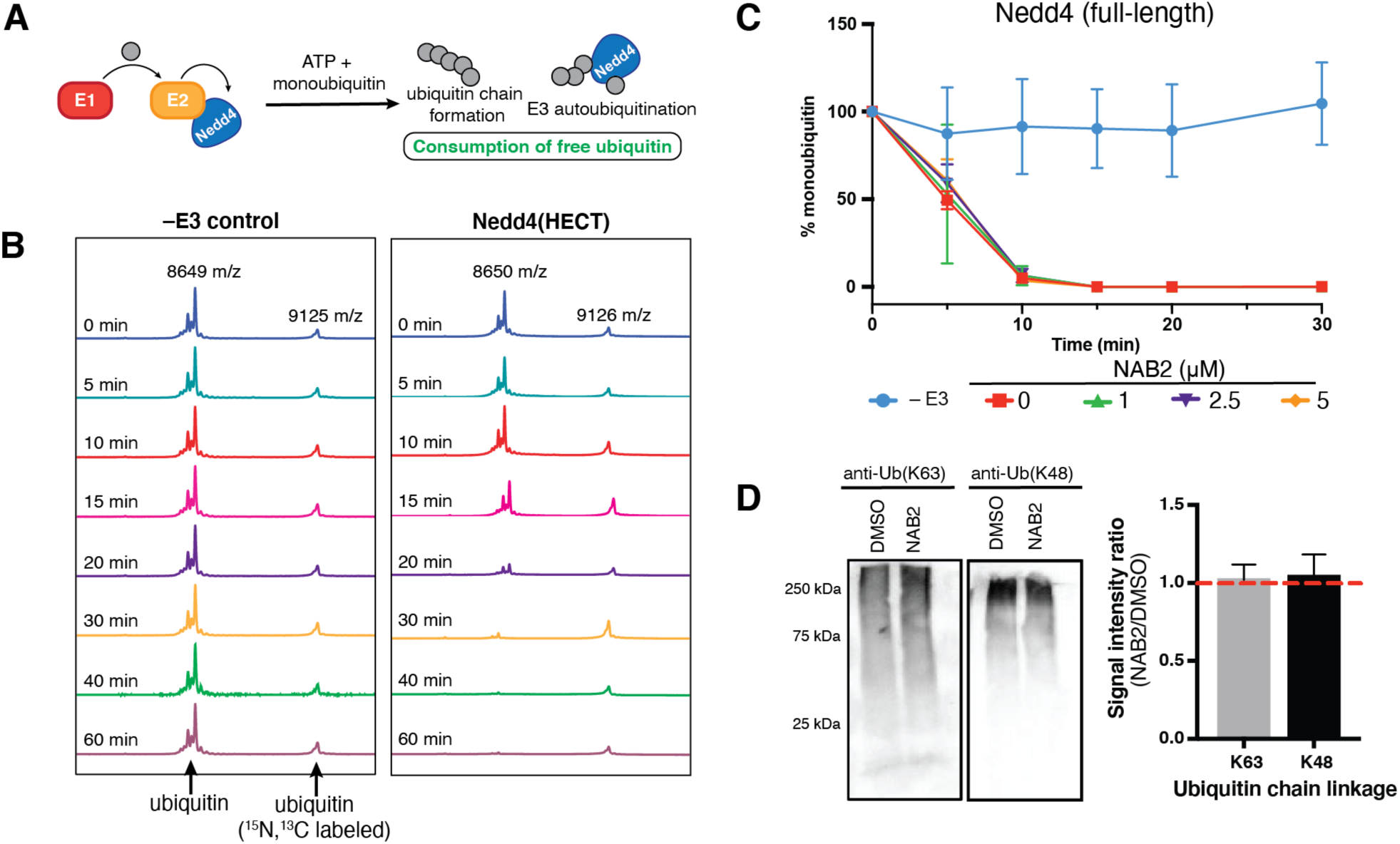
NAB2 treatment does not alter Nedd4 activity or conformation. **(A)** *In vitro* analysis of E3 ligase activity can be conducted in the absence of substrate wherein ligase activity results in formation of ubiquitin chains or E3 ligase autoubiquitination. Consumption of monoubiquitin can be monitored by MALDI-TOF and can be quantified by normalization to an internal standard. **(B)** Representative MALDI-TOF traces show time-dependent monoubiquitin consumption by Nedd4(HECT). Assays were conducted in the presence of E1 (UBE1, 50 nM), E2 (Ubch5a, 250 nM) and E3 (Nedd4(HECT), 500 nM) with ATP (1 mM) and were initiated with the addition of ubiquitin (10 *µ*M). Samples (5 *µ*L) were taken at indicated time points and quenched with 10% TFA (2 *µ*L). For MALDI-TOF analysis, samples were mixed with DHAP matrix and ^15^N,^13^C-ubiquitin (4 *µ*M) as an internal standard in a 2:1:3 ratio (sample:standard:matrix). The results of this assay show that ubiquitin signal decreases over time in the presence of Nedd4(HECT) relative to a no-E3 control. **(C)** For quantification, ubiquitin signal is normalized to ^15^N,^13^C-ubiquitin that is doped into samples after quenching and prior to MALDI-TOF analysis. Quantitative MALDI-TOF analysis shows no significant change in the rate of full-length Nedd4-dependent monoubiquitin consumption in the presence of NAB2 relative to a DMSO control. Data shown as average of triplicate ± s.e.m. **(D)** Endpoint *in vitro* assays of Nedd4 activity allow for immunoblotting-based detection of Nedd4-dependent ubiquitin chain linkage specificity. Full-length Nedd4 (5 *µ*M) was incubated with UBE1 (100 nM), Ubch5a (1 *µ*M), ubiquitin (100 *µ*M) and NAB2 (20 *µ*M), and reactions were initiated with the addition of ATP (2 mM). After 60 minutes, reactions were quenched with SDS-PAGE loading buffer and analyzed by SDS-PAGE coupled immunoblotting with ubiquitin linkage specific antibodies. The signal intensity was quantified by ImageJ and relative chain formation was measured as the intensity ratio in NAB2-versus DMSO-treated samples. Nedd4 produces K48- and K63-linked chains as its primary products, and the *in vitro* specificity of chain linkage is not altered by NAB2 treatment relative to DMSO control. A representative blot is provided and data is shown as average of triplicate ± s.e.m.

We next interrogated the linkage specificity of ubiquitin chains assembled by Nedd4. Nedd4 has been previously shown to assemble K48- and K63-linked ubiquitin chains as its primary ubiquitination products.^33^ As the identity of ubiquitin chain linkage dictates the physiological fate of the ubiquitinated substrate,^34,35^ we sought to determine if NAB2 treatment altered Nedd4 linkage specificity and could alter Nedd4-dependent polyubiquitin chain specificity. Using an *in vitro* ubiquitination assay, we detected no difference in the formation of K48- or K63-linked chains after NAB2 treatment relative to DMSO control (Supplemental Figure 3; Figure 3D). The lack of NAB2-dependent alteration of Nedd4 kinetics or ubiquitin linkage specificity *in vitro* suggested that a cellular model would be needed to uncover a potential mechanism of action.

### Confocal immunofluorescence microscopy to map NAB2-dependent changes in Nedd4 localization

Nedd4 activity is tightly regulated^23–26^ and is dependent upon protein-protein interactions with its upstream E2 conjugating enzyme and downstream substrate. Since NAB2 did not affect Nedd4 activity, we hypothesized that instead it may recue interactions between Nedd4 and proteins involved in trafficking and transport (processes disrupted by α-synuclein toxicity). We first used subcellular fractionation to quantify the distribution of Nedd4 across compartments and determined that NAB2 treatment does not alter the subcellular compartmentalization of Nedd4 (Supplemental Figure 4). Alternatively, we used confocal microscopy to visualize the co-localization of Nedd4 with Rab5a, GLG1, and calreticulin as markers of endosomes, Golgi apparatus, and endoplasmic reticulum, respectively (Figure 4B). The confocal analysis suggested that Nedd4 exhibits moderate co-localization with Rab5a, GLG1 and calreticulin in untreated samples (Figure 4; as indicated by a positive Pearson’s correlation coefficient, PCC). Comparison of PCC values for NAB2-treated samples showed that the co-localization of Nedd4 with Rab5a, GLG1, and calreticulin increased slightly after NAB2 treatment, with only Rab5a showing a significant change in co-localization (Figure 4B). These results indicate that NAB2 treatment does not significantly alter the co-localization of Nedd4 with the ER and Golgi despite the fact that NAB2 rescues ER-to-Golgi transport,^1^ but it does provide insight into the potential proximity-driven specificity of Nedd4 as a regulator of trafficking processes, as the protein exhibits moderate co-localization with the ER and Golgi apparatus prior to NAB2 treatment.

**Figure 4.**
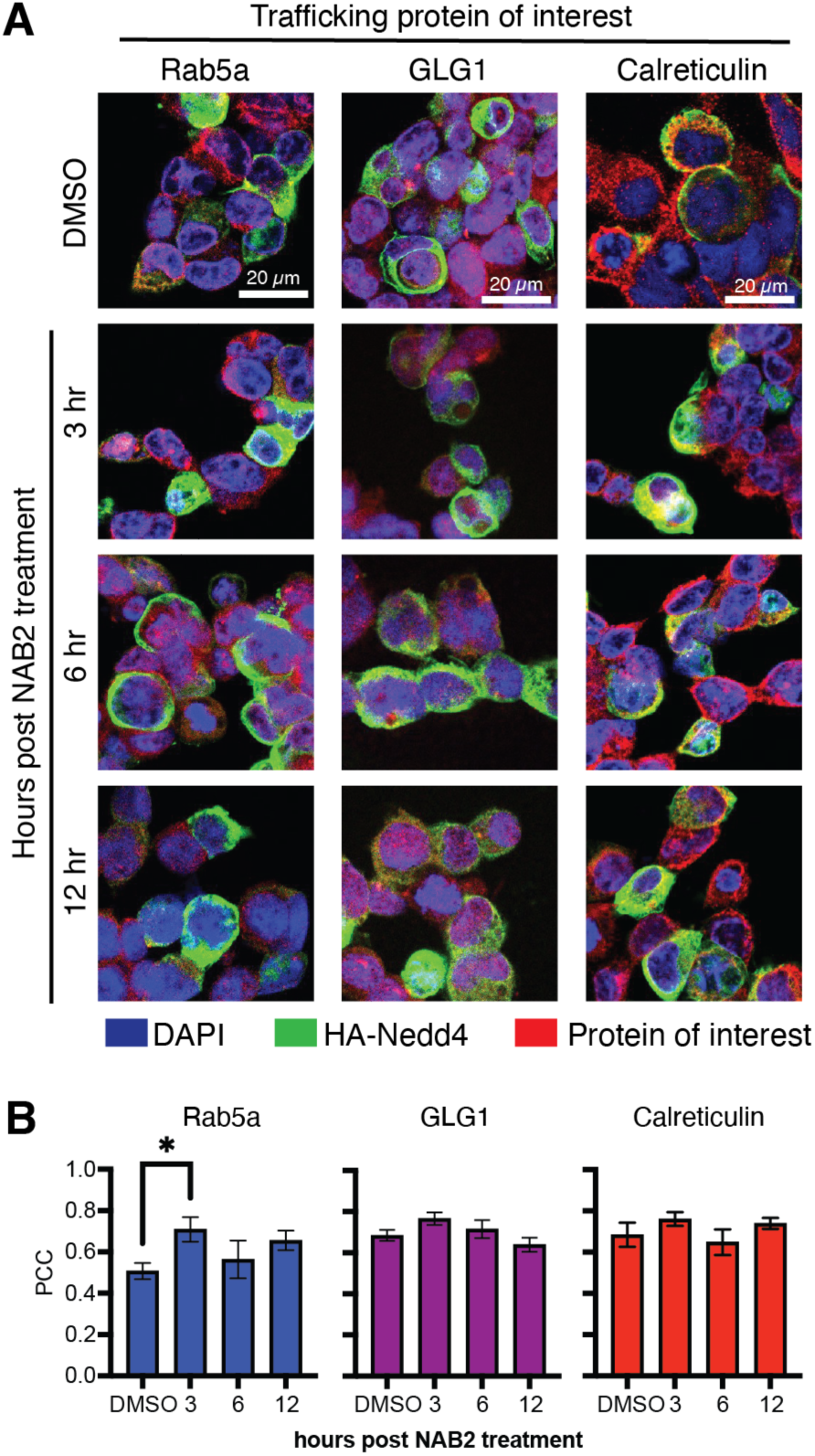
Localization experiments enable measurement of Nedd4 localization and co-localization with trafficking organelles. Confocal immunofluorescence microscopy shows moderate co-localization of HA-Nedd4 with Rab5a, GLG1, and calreticulin, well-established protein markers of the endosome, Golgi, and endoplasmic reticulum, respectively. Time-dependent NAB2 treatment shows slight increases in Nedd4 localization with the trafficking organelle markers with a significant change only present at 3 hr post NAB2 treatment in the Rab5a sample. Co-localization was measured volumetrically in single cells (defined as regions of interest, ROI) and quantified using Pearson’s correlation coefficient (PCC). Representative images **(A)** are shown and PCC data **(B)** is shown as average PCC ± s.e.m. of a minimum of 15 single cells across three microscopy frames. Statistical significance was calculated with an unpaired t-test comparing h.p.t. samples to DMSO control. For all samples, there is no significant difference unless specifically denoted where * = p < 0.05. Image analysis was conducted with Imaris and data analysis in Prism (GraphPad).

### Ubiquitin enrichment-coupled proteomic analyses for assessment of NAB2-dependent changes in the ubiquitylome

As NAB2 did not alter Nedd4 activity *in vitro* or Nedd4 co-localization with the ER/Golgi system, we hypothesized that it may instead alter the target specificity of Nedd4, which could be reflected by changes in the global ubiquitylome. As a proof-of-concept screen, we employed a tandem ubiquitin binding entity (TUBE) enrichment approach coupled to qualitative liquid chromatography-tandem mass spectrometry (LC-MS/MS) to investigate the ubiquitinome in response to time-dependent NAB2-treatment versus a DMSO control (Figure 5A). Specifically, neuroblastoma-derived SHSY5Y cells were treated with either NAB2 (20 *µ*M) for 6–24 h versus DMSO control, and the ubiquitylome was enriched by a pan-specific K48/K63-linked ubiquitin TUBE pulldown followed by LC-MS/MS.

**Figure 5.**
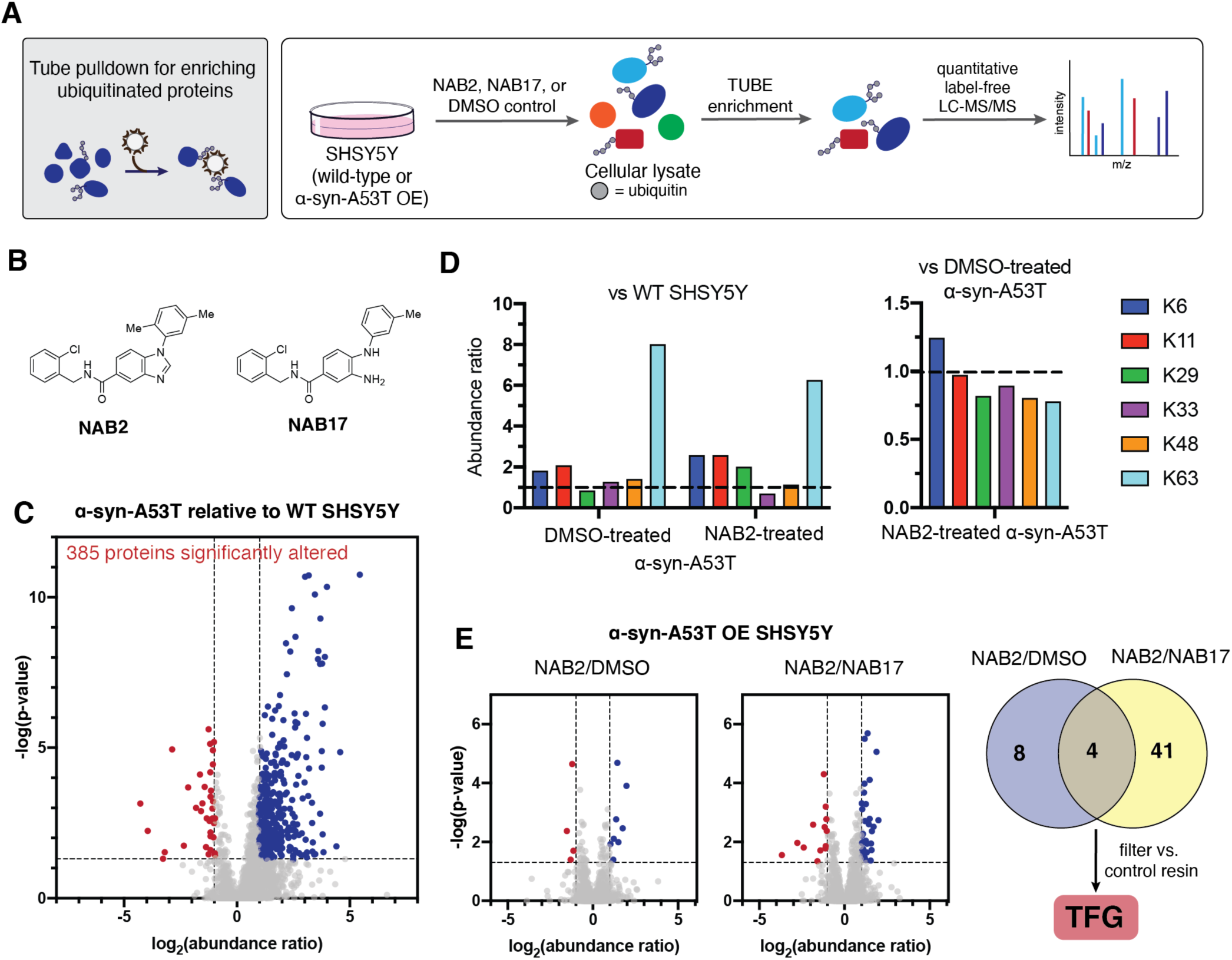
Proteomic analysis of the global ubiquitylome reveals α-synuclein- and NAB2-dependent changes in trafficking and transport protein ubiquitination. **(A)** Ubiquitinated proteins were enriched from wild-type (WT) or α-synuclein-A53T overexpressing (α-syn-A53T OE) SHSY5Y cells treated with NAB2 (20 *µ*M), NAB17 (20 *µ*M) or DMSO control. Following treatment, cells were harvested and lysed and ubiquitinated proteins were enriched by pan-specific Tandem Ubiquitin Binding Entity (TUBE) pulldown. Following enrichment, ubiquitinated proteins characterized by proteolysis-coupled LC-MS/MS and quantitative abundance changes were measured across three replicates with label free quantitation. **(B)** NAB2 was identified as the highest activity derivative while NAB17 was phenotypically inactive^1^ and was used as a second control in our proteomic analyses. **(C)** α-Synuclein overexpression induces dramatic changes in the ubiquitylome with 385 proteins showing significant alterations in abundance relative to wild-type SHSY5Y cells (significance deterimined using ± 2-fold change and p<0.05 cutoffs). **(D)** α-synuclein overexpression alters the ubiquitylome at the level of ubiquitin linkage prevalence, with up to 8-fold changes in linkage abundance in the presence of toxicity relative to wild-type cells. NAB2 treatment decreases the ratio of K63 linkage relative to untreated α-syn-A53T OE. **(E)** NAB2 treatment of (α-syn-A53T OE) SHSY5Y cells provides 12 and 45 significant hits relative to DMSO and NAB17 controls, respectively, with 4 hits identified across both control samples. Further filtering by comparison to a negative control resin pulldown reveals one remaining hit protein, TFG.

Qualitative proteomic characterization of NAB2-dependent changes in the total ubiquitylome identified 2841 total proteins across all samples (Supplemental Figure 5A). Within the total pool of identified proteins, 532 unique, non-redundant proteins were found to be enriched in the ubiquitylome only after NAB2 treatment (across all time points). This subset of proteins qualitatively indicates that NAB2 treatment alters the ubiquitylome and supports our hypothesis that NAB2-induced changes in Nedd4 specificity could be reflected at the level of the global ubiquitylome. Interestingly, functional annotation of the enriched ubiquitylome via the PANTHER^36,37^ classification system indicates that the proteins present in the NAB2-dependent sub-ubiquitylome contain a higher proportion of trafficking and transport associated proteins compared to the total ubiquitylome or DMSO control (Supplemental Figure 5B,C). The qualitative analysis thus provided proof-of-concept results that informed the design of subsequent quantitative experiments.

To better model the pathophysiological conditions that are rescued by NAB2, we applied this approach to quantify changes in protein ubiquitination in a NAB2-dependent manner. To this end, we conducted quantitative analyses of TUBE-enriched ubiquitylomes of both wild-type SHSY5Y cells and in SHSY5Y cells overexpressing A53T α-synuclein, the mutant form of α-synuclein that has been shown to induce protein aggregation and engenders trafficking defects.^6^ We also included an additional control, NAB17, a NAB analogue identified in the original SAR study as a phenotypically inactive derivative (Figure 5B).^1^ For the quantitative proteomics experiments, SHSY5Y (WT or α-synuclein-A53T overexpressing) cells were subsequently treated with DMSO, NAB2, or NAB17 for 12 hours followed by quantitative ubiquitylome analysis using TUBE enrichment and label-free LC-MS/MS (Figure 5A). In total, 2,893 proteins were quantified across the six conditions; of these, 2303 proteins were more confidently quantified by 2 or more unique peptides.

We anticipated that α-synuclein toxicity would significantly alter the ubiquitylome as part of the cellular response to the stress stimulus and that the effect of NAB2 on ER-to-Golgi trafficking may be differ in the presence of α-synuclein toxicity relative to wild-type cells. To test these hypotheses, we first compared the effect of α-synuclein-A53T overexpression on the ubiquitylome relative to wild-type cells (Figure 5C). Using a ± 2-fold change and p<0.05 cutoffs, 385 proteins in the ubiquitylome (∼17% of the total 2,303 quantified) were significantly altered in cells expressing the toxic mutant of α-synuclein relative to wild-type cells. Functional annotation and GO enrichment using the GOrilla algorithm^38^ shows α-synuclein toxicity enriches ubiquitination of proteins associated with gene expression, protein synthesis, and metabolic processes relative to the total ubiquitylome (Supplemental Figure 6). These results are consistent with previous characterizations of processes disrupted by α-synuclein-associated ER and cellular stress.^39–45^ Further analysis of ubiquitin linkage abundances (determined by comparison of diagnostic GG-K peptide abundances between samples) revealed that induction of α-synuclein toxicity increases the amount of K6, K11, K33, K48, and K63 linked ubiquitin chains relative to wild-type controls (Figure 5D). Strikingly, K63 linkages, which primarily trigger endocytosis and lysosomal degradation of plasma membrane proteins,^46^ are enriched 8-fold relative to the wild-type sample. Nedd4 assembles K63 chains as a primary product^47^ and has been well-established to be involved in the response to α-synuclein toxicity; thus, it is feasible that Nedd4 is involved in the increase in K63 chains observed in this investigation.

To further investigate potential role of Nedd4 in the response to α-synuclein induced changes in the ubiquitylome, we compiled a Nedd4 interactome from various protein-protein interaction databases (see Methods for details) for comparison with our proteomics dataset. Cross-reference of the total quantitative proteomics dataset with the compiled Nedd4 interactome reveals 104 previously identified Nedd4 interactors, while the set of proteins significantly altered by α-synuclein toxicity contains only 5 previously annotated interactors (Supplemental Table 3). This result indicates that the Nedd4 interactome is potentially largely underexplored and unannotated, particularly in regard to how its interactions are altered by cellular stimuli like α-synuclein toxicity. The effect of cellular stress on Nedd4 specificity will be explored in future investigations.

To complement the *in vitro* analyses of the NAB2 mechanism described above, we first analyzed the effect of NAB2 treatment on GG-K peptide abundances (Figure 5D). NAB2-treated α-Syn-A53T OE cells show a decrease in K63 ubiquitination relative to DMSO-treated cells, indicating that there is a NAB2-dependent shift in the ubiquitylome, but NAB2 treatment in this experiment does not fully restore cells to the ubiquitinated peptide abundances observed in wild-type cells. To identify specific proteins potentially involved in the NAB2-dependent network, we next compared the effect of NAB2 treatment in α-synuclein toxic cells (α-Syn-A53T OE SHSY5Y) relative to DMSO- and NAB17-treated controls (Figure 5E). Comparison of NAB2-treated samples shows significant alteration of 12 and 45 proteins in the ubiquitylome relative to the DMSO and NAB17 controls, respectively. Cross-reference of the 12 and 45 significant hits identified by comparison to DMSO- and NAB17-control samples reveals four proteins identified in both control experiments (Figure 5E, right). These hits include aryl hydrocarbon receptor (AHR), myosin-14 (MYH14), myosin-9 (MYH9) and protein TFG (TFG). To ensure that the identified hits were enriched with high specificity during the TUBE pulldown, we further filtered by comparison of abundance ratios in DMSO-treated α-Syn-A53T OE SHSY5Y relative to a negative control resin pulldown (using ± 2-fold change and p<0.05 cutoffs). With this additional filter, TFG is the only remaining hit protein. TFG has been previously reported as a substrate of ubiquitination,^48–54^ so to better understand if it was a potential Nedd4 substrate, we turned to bioinformatic analyses of its primary protein sequence and interactomes. Nedd4 contains four WW domains which recognize substrates that contain proline-rich PY motifs (e.g. PPxY, LPxY). TFG contains several proline-proline dipeptides and a QPPY sequence but does not contain a canonical PY motif. Further, analysis of the Nedd4 interactome reveals that TFG has not been annotated as a Nedd4 interactor, though we have noted that the Nedd4 interactome may be largely underexplored. While further interrogation of Nedd4 interactions with TFG and of its larger interactome is needed, it is outside the scope of this investigation and will be explored in future analyses.

With the role of Nedd4 and NAB2 treatment in rescuing α-synuclein associated trafficking defects in mind, we sought to further understand the potential role of TFG in this process through functional and interaction network analyses. Excitingly, TFG is involved in the regulation of ER-to-Golgi trafficking and transport,^55–57^ the process that is disrupted by α-synuclein toxicity and was shown to be rescued by NAB2 treatment. To further understand the possible connections between TFG, Nedd4, and α-synuclein at a cellular level, we generated a merged interaction network of the experimentally annotated interactome of each of these proteins using the IntAct^58^ and BioGrid^59^ protein-protein interaction databases (Figure 6A). Inclusion of Nedd4 and α-synuclein in the merged interaction network enables identification of putative functional connections between the NAB2-dependent hit and the proteins involved in the toxicity process or rescue thereof. Interestingly, there are several proteins that are shared between the interactomes of Nedd4, α-synuclein, and TFG (Figure 6). There are thirteen and seven interactors shared between TFG and Nedd4 or α-synuclein, respectively, and there are four common interactors shared between all three proteins of interest (Nedd4, α-synuclein, and TFG; Figure 6B). Functional annotation reveals that 13 of the 24 shared interactors are associated with trafficking and transport processes, establishing a biological pathway as a link the NAB2-dependent hit TFG and Nedd4 and α-synuclein. Of particular interest in the shared interactome are proteins such as LRRK2 (a kinase that is well-established for its role in a genetic form of Parkinson’s disease),^60^ GABARAP, GABARAPL1/GABRAPL2, as well as MAP1LC3A and MAP1LC3B (called LC3s). The latter five proteins are key regulators in autophagosome formation and maturation, and maintenance of cellular homeostasis.^61^ While we anticipate that other proteins, including those outside of the enriched ubiquitylome, may be involved in the NAB2-dependent rescue of ER-to-Golgi trafficking, the use of quantitative TUBE-coupled proteomics provides insight into the effect of both α-synuclein toxicity and NAB2 treatment on the ubiquitylome through identification of global α-synuclein dependent alterations and identification of trafficking associated protein TFG that is modulated in a NAB2-dependent manner. Further, bioinformatic analysis complements the proteomic results by revealing a putative protein network connecting the proteins of interest.

**Figure 6.**
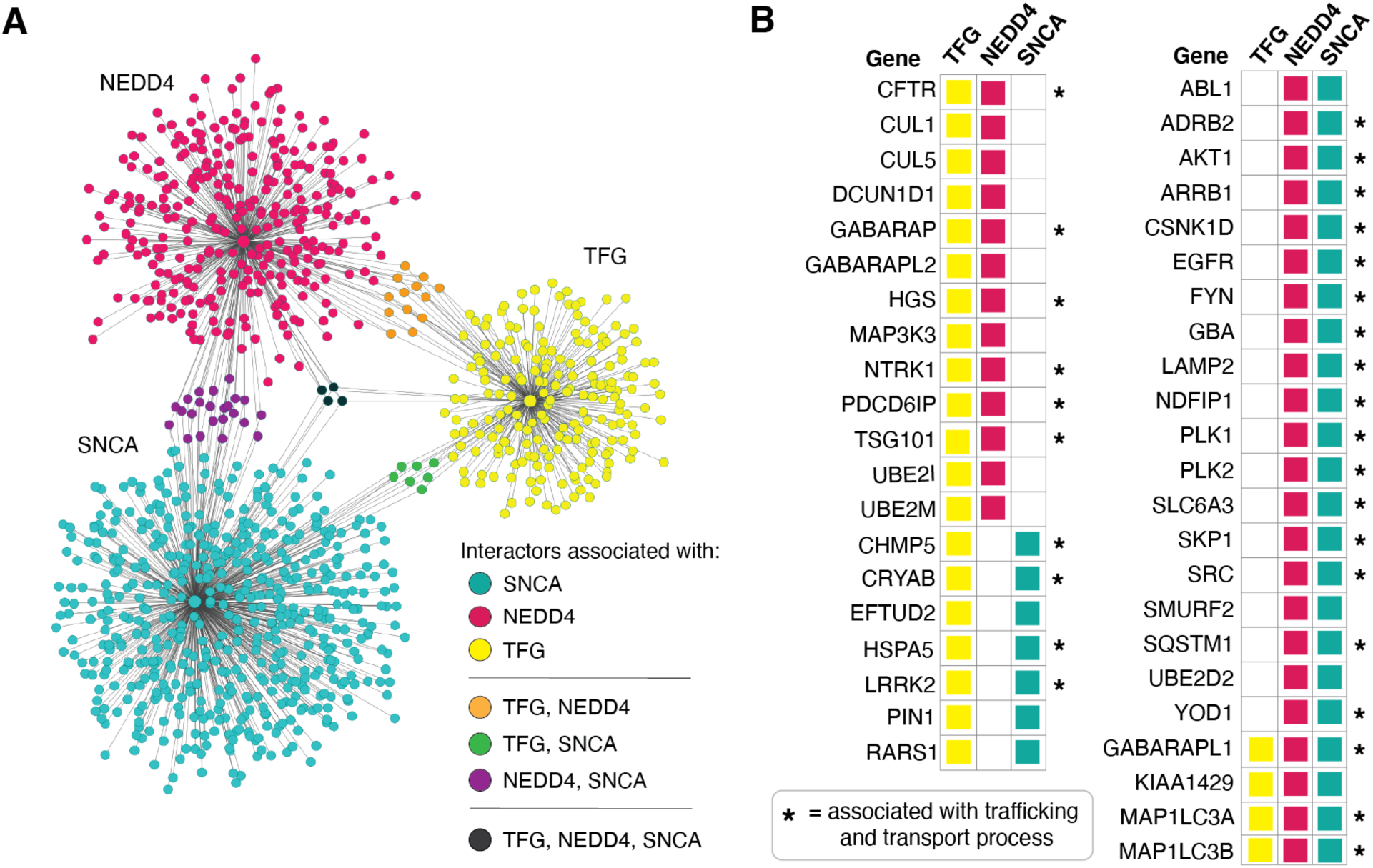
Analysis of hits identified in quantitative identification of NAB2-dependent ubiquitylome. **(A)** Merged protein-protein interaction networks from the IntAct and BioGrid databases show interactors of the significant hit from NAB2 treatment of α-synuclein toxic SHSY5Y cells (α-synuclein-A53T overexpression, Figure 5E and Table 1). Networks visualized using Cytoscape. Merging networks for hits and α-synuclein as well as Nedd4 enable identification of interactors shared between **(B)** significant TFG as well as proteins that are shared interactors between TFG and α-synuclein and Nedd4. The networks in which each shared protein is found are indicated by color blocks which correspond with the legend in **(A)**. Asterisk (*) indicates a protein that is functionally annotated (GO annotation) as a trafficking and transport associated protein.

## Discussion

Initial identification of NAB2 as a potential anti-Parkinsonian lead compound showed its activity was dependent upon E3 ubiquitin ligase Nedd4,^1^ but target engagement and biochemical validation of the NAB2 mechanism of action was unexplored. Herein, we present a biochemical and biophysical interrogation of NAB2 target engagement with Nedd4, a putative target identified through chemical genetic screening. By SPR, ITC and PTSA, we observed that NAB2 bound to Nedd4 with nM affinity (K_D_^*app*^), which was localized to an N-terminal region of Nedd4 that lied outside of the C-terminal catalytic HECT domain. Despite evidence that NAB2 binding does not occur in the catalytic domain of the ubiquitin ligase, we sought to characterize the enzymology of Nedd4 in a NAB2-dependent manner as the ligase activity is tightly regulated by interactions between its catalytic domain and upstream structural components (C2 domain, WW domains, and linker regions).^23–26^ *In vitro* analyses of Nedd4 activity reveal that NAB2 does not alter the global conformation, activity, or ubiquitin linkage specificity of Nedd4 as measured by bottom-up HDX-MS and quantitative activity assays. While HDX-MS results indicate that NAB2 does not induce a global conformational change, there are small regions in which deuterium uptake changes in a NAB2 dependent manner. These areas are distributed across the N-terminal end of the protein upstream of the catalytic HECT domain, a result which is consistent with our biophysical and thermodynamic measurements of NAB2 binding in which NAB2 binds to full-length Nedd4 but not the isolated HECT domain. The other *in vitro* analyses of Nedd4 activity (measured via MALDI-TOF quantification of ubiquitin consumption rate) and ubiquitin linkage specificity indicate no global NAB2-dependent change in the backbone conformation, but is consistent with a side-chain mediated interaction that doesn’t inhibit its E3 ligase activity, but most likely modulates its substrate specificity *in vitro*. It is important to note, however, that Nedd4 has many possible interactions with upstream E2 conjugating enzymes and downstream substrates, which are likely regulated by cellular stimuli. It is not sufficient, therefore, to only consider the results of *in vitro* analyses with limited interactors. We therefore complemented the *in vitro* analyses with cellular and proteomic analyses to better understand the mechanism of NAB2 and role of Nedd4 at a more global level.

As NAB2 was initially shown to rescue ER-to-Golgi trafficking defects in a Nedd4-dependent manner, we turned to cellular and proteomic analyses to identify NAB2-dependent changes in Nedd4 localization with cellular trafficking machinery. Confocal microscopy measurement of Nedd4 co-localization with markers of trafficking organelles shows that Nedd4 exhibits a moderate co-localization with the Golgi and ER at basal conditions and that NAB2 treatment induces slight but insignificant increases in Nedd4 localization with these organelles. Interestingly, NAB2 treatment induces a significant increase in Nedd4 co-localization with Rab5a, a marker of the early endosome. Nedd4 has been previously implicated in endosomal trafficking of α-synuclein through K63 linked ubiquitin chain assembly, but there is not previous evidence of NAB2-dependent Nedd4 involvement with endosomal trafficking. We anticipate that the role of Nedd4 and NAB2 in endosomal trafficking could be explored more broadly in future investigations. Subsequent proteomic interrogation of the ubiquitylome using TUBE-coupled proteomics indicates that α-synuclein toxicity drastically remodels the ubiquitylome (both at a protein level and by changing the prevalence of specific ubiquitin linkage types), and functional analysis reveals that the significantly altered proteins are involved in processes previously reported to be disrupted by α-synuclein. Further analysis reveals a NAB2-dependent decrease in K63 ubiquitin chain linkage abundances relative to untreated α-synuclein toxic cells.

Analysis of NAB2-treated α-synuclein toxic SHSY5Y cells revealed four significant hits relative to DMSO- and NAB17-treated controls. These hits include: AHR, MYH9, MYH14, and TFG. Of these hits, three (MYH9, MYH14, and AHR) did not vary significantly from control resin pulldowns, leaving TFG as the only remaining hit. TFG has been established as a regulator of ER-to-Golgi trafficking, the process that is disrupted by α-synuclein toxicity and restored by NAB2 treatment. While TFG has not been previously annotated as a Nedd4 substrate, bioinformatic analysis of the TFG, α-synuclein, and Nedd4 interactome reveal a network of functionally related, shared interactors and establishes a putative network of trafficking and transport proteins that link the NAB2-dependent hit with Nedd4 and α-synuclein. Further, cross-reference of the Nedd4 interactome with proteomic hits shows that few proteins in the α-synuclein- or NAB2-dependent ubiquitylome are known Nedd4 interactors, indicating that the Nedd4 interactome is largely unexplored. Thus, further analysis of Nedd4 interactions in the cell would provide insight into the role of the ligase in the rescue of α-synuclein toxicity. Despite this, the proteomics results provide a putative network of trafficking associated proteins that are altered in the ubiquitylome in a NAB2-dependent manner.

Cumulatively, the analyses described herein provide insight into the effects of α-synuclein toxicity and NAB2 treatment on the ubiquitylome and demonstrate that NAB2 binds to Nedd4 without altering its enzymatic activity or conformation *in vitro* and largely does not alter Nedd4 cellular localization relative to trafficking organelles. Thus, these results indicate that NAB2 binding may induce a phenotypic change via a more nuanced or complex mechanism, and our proteomic characterization sheds light on proteins potentially involved in NAB2-dependent restoration of trafficking defects. The efforts described highlight the power of phenotypic screens for the identification of novel ligands that can alter complex biological processes (like those associated with neurodegeneration)^62^ but also demonstrate that ligands identified in this manner may act via mechanisms that are not easily unraveled. While the use of these phenotype-driven platforms enables identification of novel ligands like NAB2, there are many opportunities that remain in fully elucidating the mechanism by which NAB2 rescues α-synuclein toxicity. The efforts toward biochemical and proteomic validation provide valuable insight into possible NAB2 mechanisms and lay the groundwork for exciting biochemical investigations of NAB2 targets, Nedd4 specificity, and α-synuclein toxicity that are currently underway.

## Significance

Limitations in the long-term efficacy of Parkinson’s therapeutics has motivated research to identify novel drug targets and lead compounds. This process typically relies upon phenotype-driven screens for identification of compounds that can alleviate complex biological defects related to the neurodegenerative disorder. While these approaches are powerful, they often result in finding lead compounds for which the specific cellular target or mechanism of action is not known. To advance the compounds in therapeutic development, there is great utility in unravelling the underlying mechanisms. Herein, we interrogate the mechanism of one such compound, NAB2, that was identified through phenotypic screening. NAB2 was shown to rescue α-synuclein associated trafficking defects, and its activity was dependent upon E3 ubiquitin ligase Nedd4.^1,13^ We demonstrate that NAB2 binds to Nedd4 with high apparent affinity but does not alter its activity or conformation *in vitro*. These biophysical and biochemical analyses are complemented by proteomic identification of NAB2-dependent changes in global protein ubiquitination through which we show that NAB2 stimulates the ubiquitination of trafficking associated proteins. Our findings provide a putative NAB2-dependent protein network involved in cellular trafficking and highlight the utility of interrogating the mechanism of ligands identified through phenotypic screens. Further, we demonstrate that the proteins affected by NAB2 are related to other Parkinson’s associated pathways, laying the groundwork for further investigation of NAB2 specificity and translatability to other neurodegeneration related trafficking defects.

## Acknowledgements

The authors kindly thank Dr. Brian Watts of the Duke Human Vaccine Institute (DHVI) for providing access to and training for Surface Plasmon Resonance experiments and Dr. Peter Silinski of the Duke Chemistry Shared Instrument Facility for assistance with implementation of the MALDI-TOF assay platform and HRMS. Additionally, the authors also thank Addgene for access to plasmids (as described in Supplemental Material) made available through the plasmid repository. The authors wish to acknowledge the Duke School of Medicine and the Duke Cancer Institute for the funding of the Lumos mass spectrometer used for proteomics analyses. Finally, the authors gratefully thank the members of the McCafferty lab for thoughtful feedback during the course of this project and in preparation of this manuscript.

## Author contributions

D.G.M. supervised the work described herein, secured funding, mentored A.K.H. and H.D.A., and aided in manuscript preparation and editing. A.K.H. participated in conceptualization, conducted all experiments, data collection and visualization, and prepared the manuscript. H.D.A. assisted in protein purification and in establishing and validating *in vitro* assays.

## Funding

This work was kindly supported by Duke University, National Institutes of Health Grant 1R21NS112927-01 to D.G.M., Michael J. Fox Foundation Grant 16250 to D.G.M., National Science Foundation Grant MCB1020649 to S.W.E., and National Science Foundation Graduate Research Fellowship GRFP 2017248946 to A.K.H.

## Materials and methods

### Plasmids

All plasmids used in the described experiments were ordered from Addgene plasmid repository (Supplemental Table 1). Plasmids were sequenced upon arrival and used as received unless specifically noted. Any modifications to plasmid sequences were generated by Q5 site-directed mutagenesis (New England BioLabs) and confirmed by sequencing. All plasmid information, including accession numbers, expression constructs, and modifications by site-directed mutagenesis, are described in Supplemental Table 1.

### Protein expression and purification

All proteins described herein for *in vitro* analyses were expressed and purified as recombinant proteins with the exception of E1 activating enzyme UBE1 and heavy isotope labeled ubiquitin (both purchased from R&D Systems). Recombinant ubiquitin, Ubch5a, Nedd4(HECT), and full-length Nedd4 were accessed by expression in *E. coli* using the expression constructs described in Supplemental Table 1. For all recombinant proteins described herein, expression and purification followed the general method described below unless otherwise stated, and full expression and purification details are described in Supplemental Methods.

The expression plasmid was transformed into the desired competent *E. coli* strain and were plated on an antibiotic selection plate for overnight growth at 37 °C. Starter cultures were inoculated from the selection plate and grown overnight (200 rpm, 37 °C). Expression cultures were inoculated from saturated starter culture and OD_600_ was monitored until the desired density at which point expression was induced by addition of isopropyl β-D-1-thiogalactopyranoside (IPTG). Following expression, cells were harvested by centrifugation (Sorvall SLA-3000, 5,000 rpm, 20 min, 4 °C, resuspended in lysis buffer, and lysed using an EmulsiFlex C5 homogenizer (Avestin). Cell debris was removed from lysates by ultracentrifugation (Beckman Coulter 70-Ti rotor, 40,000 rpm, 4 °C, 0.02 Torr, 45 minutes) and lysates were purified by affinity chromatography via AKTA FPLC (GE Life Sciences).

### Mammalian tissue culture

SHSY5Y cells were obtained from the Duke University Cell Culture Facility and were maintained at 37 °C with 5% CO_2_ in DMEM:F12 (1:1; Gibco) supplemented with 10% FBS (Sigma Aldrich) and 1X penicillin-streptomycin (Gibco). Cells were passaged at 80% confluency with a sub-cultivation ratio of 1:250 unless seeding for transfection experiments where a higher density was required. Transfection of SHSY5Y was performed with GeneX Plus (ATCC). Transfected plasmids are described in Supplemental Table 1.

### Surface Plasmon Resonance for Low Molecular Weight Kinetics

NAB2 binding to GST-Nedd4 and Nedd4(HECT) was characterized via SPR using the Biacore T200 instrument (GE Life Sciences). Recombinant protein was purified as described with an additional purification step using size exclusion chromatography to ensure high purity prior to use in SPR. SPR analyses were performed with sterile filtered, degassed PBS. Protein was immobilized to a Series S CM5 chip using NHS crosslinking to a surface RU of 7,000-9,000 to provide high density for low molecular weight kinetics. Following immobilization and equilibration, the protein surface was exposed to a concentration gradient of NAB2 (0 to 100 *µ*M in PBS with <1% DMSO for solubility). Binding was calculated with 1:1 binding algorithm for low molecular weight kinetics.

### Protein thermal shift assay for measurement of ligand-induced protein stability changes

Nedd4(HECT) or full-length Nedd4 (4 *µ*M) was incubated with NAB2 at various concentrations for 30-60 minutes. Following incubation, SYPRO orange (5000X stock in DMSO, Invitrogen) was added to a final concentration of 5X and mixtures were aliquoted to a final volume of 15 *µ*L into a LightCycle 96-well white qPCR plate (Roche). Thermal denaturation was conducted in a LightCycler 480 (Roche) via continuous heating from 20 to 85 °C over 18 minutes. Experimental T_m_ was calculated as the absolute minimum of the negative first derivative curve of the melting curve (-d(fluorescence intensity)/d(temperature)).

### Ubiquitination activity assay by immunoblotting

*In vitro* ubiquitination activity assays were conducted as endpoint assays via immunoblot detection using recombinant E1, E2, and E3 enzymes. E1 (100 nM), E2 (1 *µ*M) and E3 (5 *µ*M) were incubated with ubiquitin (100 *µ*M) in reaction buffer (100 mM Tris, 25 mM MgCl_2,_ 0.1% Tween, pH 8) and reaction was initiated by addition of ATP (2 mM). Reactions were incubated for 1 hour at 37 °C and quenched with the addition of SDS-PAGE loading buffer. Samples were heated to 95 °C for 10 minutes and separated by SDS-PAGE. Gels were transferred to PVDF membrane for immunoblotting and ubiquitination activity was detected by blotting with anti-ubiquitin (1:2000, Abcam ab7780) or linkage specific antibodies (Abcam: anti-ub(K63), ab179434; anti-ub(K48), 140601) and HRP-conjugated secondary antibody (BioRad). Signal was detected with ECL reagents (Genesee Scientific) and quantified using ImageJ.

### MALDI-TOF ubiquitin activity assay

Ubiquitination activity was quantified in a time-dependent manner by MALDI-TOF detection of monoubiquitin consumption according to a procedure adapted from De Cesare et al.^63^ E1 (50 nM), E2 (250 nM), and E3 (500 nM) were combined in reaction buffer composed of 0.25 mg/mL BSA in 10 mM HEPES pH 8.5, 10 mM MgCl_2_ and 1 mM ATP. Reactions were incubated at 37 °C and initiated with the addition of ubiquitin (10 *µ*M). At desired time intervals, an aliquot (5 *µ*L) was quenched with 10% TFA (1 *µ*L). Following collection of all time points, samples were doped with DHAP matrix and 4 *µ*M ^15^N,^13^C-ubiquitin as an internal standard (3:1:2 matrix:standard:sample) and spotted on an AnchorChip 384 BC plate (Bruker Daltonics). Samples were analyzed by MALDI-TOF (Bruker Autoflex Speed LRF MALDI-TOF System) using the following automated AutoXecute method: Reflector Positive mode with laser intensity at 80%, Laser Fuzzy Control switched off, and accumulation parameters set to 4000 satisfactory shots in 500 shot steps with movement parameters set to “Walk on Spot”. Spectra were accumulated by FlexControl software and processed using FlexAnalysis software. Monoubiquitin signal was normalized to signal from the heavy isotope derivative and plotted as normalized intensity versus time to determine time-dependent ubiquitination activity of the signaling cascade.

### Hydrogen-deuterium exchange mass spectrometry (HDX-MS)

Full-length recombinant Nedd4 (50 *µ*M) was pre-incubated with NAB2 or DMSO control and then exposed to deuterated buffer to HDX analysis at time points ranging from 10 sec to 3 hr. Following exchange, the reaction was quenched to pH 2.4 with cooled quenching buffer (4 °C) to prevent back exchange of deuterium, and the sample was submitted to sequential pepsin digestion and desalting via a C18 column (at 4 °C) in line with liquid chromatography coupled tandem mass spectrometry (LC-MS/MS) for peptide identification. Peptide coverage and identity were determined using SEQUEST software with the Nedd4 sequence as reference. Relative deuterium uptake of each peptide was determined by comparison to the non-deuterated (“all hydrogen”) control using the ExMS2 program,^64^ and NAB2-dependent changes in the rate of deuterium uptake were determined by comparison of the NAB2-treated sample relative to the DMSO control. Heat maps were replotted using open source Morpheus program (Broad Institute) and relative deuterium uptake was compared between NAB2-treated and DMSO control.

### Qualitative proteomic analysis of TUBE-enriched ubiquitylome

SHSY5Y cells were seeded in 150 mm tissue culture plates and grown in DMEM/F12 (1:1) supplemented with 10% FBS and penicillin/streptomycin (1X). When cells were 80% confluent, a treatment time course was initiated. Cells were treated for 6, 12, 18, and 24 hours with 20 *µ*M NAB2 in DMSO (and a DMSO control at 24 hour timepoint). Following treatment, cells were harvested and resuspended in TUBE lysis buffer (50mM Tris-HCl, pH 7.5, 0.15M NaCl, 1mM EDTA, 1% NP-40, 10% glycerol, published by LifeSensors, Inc.) with 1X protease inhibitor cocktail (Bimake) and 50 *µ*M PR-619 (non-specific DUB inhibitor, Sigma). Cells were then sonicated (Fisher Scientific Model 120 Sonic Dismembrator; 4 pulses, 5 seconds per pulse, 30% amplitude) and cell debris was collected by centrifugation (14,000 rpm, 4 °C, 10 minutes). Lysate concentration was determined by Bradford assay. Magnetic TUBE 1 beads (pan-selective) were equilibrated by washing with TBS-T and lysate was added to TUBE beads to a final ratio of 100 *µ*L bead slurry to 1 mg total protein. An additional sample was prepared with magnetic control beads (LifeSensors) and DMSO-treated lysate. Lysate was incubated with TUBE beads or control beads for 2 hours at 4 °C with end-over-end rotation. Following incubation, beads were collected and supernatant was removed. Beads were washed with TBST and bound protein was eluted for proteomic analysis with 50 mM TEAB buffer containing 5% SDS. Subsequently, a BCA assay was performed on the samples. The control pulldown had ∼0.3 *µ*g/*µ*l and the other samples were 0.55, 0.52, 0.59, 0.5 and 0.53 *µ*g/*µ*l (for DMSO-treated, 6hr, 12hr, 18hr, and 24 hr timepoints, respectively). 25 *µ*L of each sample was reduced and alkylated, followed by clean up and digestion with trypsin using an S-Trap (Protifi). After lyophilization, tryptic digests were resuspended in 12 *µ*L, and 1 *µ*L of each sample was analyzed by LC-MS/MS (see below) followed by database searching using Mascot. Search results were annotated at a 1% peptide/protein FDR in Scaffold.

### Quantitative proteomic analyses of TUBE-enriched ubiquitylome

Fifty microliters of eluents were diluted to 75 μL with 10% SDS in 50 mM triethylammonium bicarbonate (TEAB) buffer. Samples were reduced by heating with 10 mM DTT at 80 °C for 10 min. Next, reduced thiols were alkylated with 25 mM iodoacetamide at room temperature for 30 min. Finally, samples were processed using an S-Trap micro device (Protifi) using the manufacturer’s protocol. Digestion of each sample was performed using 1 μg of Sequencing Grade Modified Trypsin (Promega) in TEAB per sample at 47 °C for 1 h. After elution from the S-Trap, peptides were lyophilized and resuspended in 12 μl of 1% TFA/ 2% MeCN. A QC pool was made by combining an equi-volume of each sample.

Quantitative one-dimensional liquid chromatography, tandem mass spectrometry (1D-LC-MS/MS) was performed on 4.5 μL of the peptide digests per sample in singlicate based on an initial loading study. After two conditioning runs with the QC pool, samples were analyzed in a batch-randomized manner with interspersed QC pools. The LC-MS/MS used a nanoACQUITY UPLC system (Waters) coupled to a Thermo Fusion Lumos high resolution accurate mass tandem mass spectrometer (Thermo) via a nanoelectrospray ionization source and FAIMS Pro Interface. Briefly the sample was first trapped on a Symmetry C18 180 μm × 20 mm trapping column (5 μl/min at 99.9/0.1 v/v H_2_O/MeCN) followed by an analytical separation using a 1.7 μm ACQUITY HSS T3 C18 75 μm × 250 mm column (Waters) with a 90 min gradient of 5 to 30% MeCN with 0.1% formic acid at a flow rate of 400 nl/min and column temperature of 55 °C. Data collection on the Fusion Lumos MS was performed in data-dependent acquisition (DDA) mode with a 120,000 resolution (@ m/z 200) full MS scan from m/z 375 to 1600, with a compensation voltage (CV) of −40, −60 or −80, and a target AGC value of 2e5 ions and 50 ms maximum injection time (IT) with internal calibration enabled. Peptides were selected for MS/MS using with advanced peak determination enabled, peptide monoisotopic peak determination, and including charge states 2-5. MS/MS used HCD fragmentation and detection in the ion trap. For each CV, a 0.66 s method used an isolation width of 1.2 m/z, a normalized collision energy of 30 ± 5%, a rapid ion trap scan rate, normalized AGC target of 100% and auto IT. A 20 s dynamic exclusion was enabled.

Following the MS analysis, data was processed using Proteome Discoverer 2.4 (Thermo). Data processing used Minora Feature Detector with min. trace length of 3, max. ΔRT of isotope patterns of 0.2 min, and PSM confidence of at least medium. Database searching was performed using Mascot 2.4 using a Swissprot database with homo sapiens taxonomy (downloaded on 031119; 20,358 unique sequences; with bovine casein appended) with trypsin specificity, up to 2 missed cleavages, precursor mass tolerance of 5 ppm, fragment mass tolerance of 0.8 Da, static carbamidomethyl(C), variable oxidation(M), deamidation(NQ) and GG(K). For label-free quantification (LFQ), Minora feature mapper used a trace length of 3 and max ΔRT of 0.2. Default percolator settings were used for FDR determination. Consensus steps used the Feature Mapper with RT alignment and a max RT shift of 5 min and min S/N threshold of 1. The precursor ions quantified used unique+razor peptides and intensity precursor abundance. Normalization was to total peptide Protein abundances were calculated using summed peptide abundances, and imputation used replacement of missing values with random values sampled from the lower five percent of detected values. Data was exported for master proteins that met a high confidence (1% peptide and protein) FDR. Statistical analysis in Proteome Discoverer used ANOVA.

The raw mass spectrometry proteomics data, the spectral library, Spectronaut .SNE file and associated results and metadata have been deposited in MassIVE (ftp://MSV000085432@massive.ucsd.edu with password “NEDD4”) with the ProteomeXchange ID PXD019245.

### Bioinformatic analysis of Nedd4 interactome and ubiquitylome

Nedd4 interactome information was compiled from various protein-protein interaction databases including BioGrid,^59^ InnateDB,^65^ MINT,^66^ Mentha,^67^ IntAct^58^ and IMEx.^68^ The compiled interactome was screened for redundancy prior to analyses. The compiled interactome was visualized by Cytoscape^69^ and was used for cross-reference to proteins identified from proteomic analyses.

Ubiquitylome proteins identified by proteomic analyses were annotated using the PANTHER functional classification system.^36^ Proteins were annotated for biological process (function) and sub-cellular compartmentalization for organelle localization.

### Subcellular fractionation and immunoblotting

SHSY5Y cells were seeded into 6-well plates and were transfected with pCl-HA-Nedd4. At 48 hpt, cells were treated with DMSO control or NAB2 (20 *µ*M) in 6 hour intervals. Following treatment, proteins were harvested from the cells by subcellular fractionation using ProteoExtract subcellular fractionation kit (Millipore Sigma). Fractionated samples were analyzed by SDS-PAGE and electrophoresis for detection of HA-tagged Nedd4 (Roche anti-HA high affinity,1:2000 dilution, catalog no. 11867423001) and actin (anti-actin, 1:2000, Abcam ab12148). Nedd4 signal was quantified by ImageJ,^70^ normalized to actin loading control signal, and plotted as percentage of total normalized HA-Nedd4 signal.

### Immunofluorescence microscopy for protein translocation experiments

SHSY5Y cells were seeded for transfection and were subsequently (24h post-seeding) transfected with pCl-HA-Nedd4 or pCDNA3.1 vector control using GeneX Plus Transfection reagent (ATCC). Forty-eight hours post-transfection, cells were treated with NAB2 (20 *µ*M) or DMSO control at 6 hour intervals. Following treatment, cells were fixed using cold methanol and blocked with 3% BSA in TBST for 30 minutes. Following blocking, cells were treated with anti-HA (Pierce High-Affinity, Roche) primary antibody and primary antibody for the trafficking marker of interest (anti-RAB5a, Fisher PA529022; anti-GLG1, Fisher PA526838; anti-calreticulin, Fisher PA3900) for one hour and then with species-specific AlexaFluor conjugated secondary antibodies and Hoechst stain. Cover slips were fixed on microscopy slides and analyzed using a Zeiss AiryScan 880 confocal microscope using the 40X oil objective. Images were processed using Imaris software. Quantitation of co-localization was determined by defining single cells expressing transfected HA-Nedd4 as regions of interest (ROI) and calculating the Pearson’s Correlation Coefficient (PCC) across the full volume (z-stack) of the ROI. A minimum of 15 ROI across three separate microscopy frames were measured for each condition and PCC was reported as average ± s.e.m. for comparison of NAB2 treated (3, 6, 12 hr post-treatment) vs time = 0 control.

## Notes

ftp://

### Competing Interest Statement

The authors have declared no competing interest.

