## Supplemental Materials for "Characterization of small molecule induced changes in Parkinson’s-related trafficking via the Nedd4 ubiquitin signaling cascade"

for

### Supplemental methods

*Protein expression and purification*

Expression construct information is detailed in Supplemental Table 1. All plasmids were ordered from Addgene and used as received unless specifically stated in the site-directed mutagenesis section below. For all expression and purification methods described below, protein expression was conducted in BL21(DE3) or BL21(DE3)-CodonPlus-RIL competent *E. coli* and was induced with the addition of isopropyl β-D-1-thiogalactopyranoside (IPTG; GoldBio) to the specified concentration. Proteins were purified by affinity chromatography as described below and purity was analyzed by SDS-PAGE and Coomassie staining. After purification and characterization, protein concentration was checked by Bradford assay and proteins were aliquoted for storage as described.

Ubch5a: pET15-Ubch5a was transformed into BL21(DE3) competent *E. coli* and successful transformants were selected for by growth on an ampicillin selection plate. For expression, a starter culture (100 mL LB media) was inoculated from a streak of colonies on the selection plate and allowed to reach saturation by overnight growth at 37 °C. Expression cultures were inoculated from saturated starter culture (10 mL starter culture per 1 L LB media) and grown at 37 °C until $OD_{600}$ = 0.6 at which time expression was inducted with the addition of IPTG to a final concentration of 100 µM and grown overnight at 18 °C. After growth, cells were harvested and pelleted for storage at –20 °C until use for purification. For purification, the cell pellet was thawed on ice and subsequent resuspended in lysis buffer (50 mM Tris, 500 mM NaCl, pH 7.5 with 1X protease inhibitor cocktail, 40 µM PMSF and lysozyme). The cells were lysed and the cell debris was removed from the lysate by ultracentrifugation. Ubch5a was purified from the supernatant of the lysate as follows: the supernatant was loaded onto a $Ni^{2+}$-charged immobilized metal affinity column (equilibrated with 50 mM Tris, 500 mM NaCl, 20 mM imidazole, pH 7.5). Ubch5a was eluted via gradient elution from 20 to 500 mM imidazole. Ubch5a-containing fractions were pooled dialyzed overnight against 4 L of dialysis buffer (50 mM Tris, 500mM NaCl, pH 7.5) to remove imidazole from the sample. The protein was subsequently concentrated and aliquoted for storage at 4 °C, –20 °C (in 40% glycerol), and –80 °C.

Nedd4(HECT): pET28-MHL-5C7J was transformed into BL21(DE3) competent *E. coli* and successful transformants were selected for by growth on a kanamycin selection plate. For expression, a starter culture (100 mL LB media) was inoculated from a streak of colonies on the selection plate and allowed to reach saturation by overnight growth at 37 °C. Expression cultures were inoculated from saturated starter culture (10 mL starter culture per 1 L LB media) and grown at 37 °C until $OD_{600}$ = 0.6 at which time expression was inducted with the addition of IPTG to a final concentration of 500 µM and grown overnight at 18 °C. After growth, cells were harvested and pelleted for storage at –20 °C until use for purification. For purification, the cell pellet was thawed on ice and subsequently resuspended in lysis buffer (50 mM Tris, 500 mM NaCl, pH 7.5 with 1X protease inhibitor cocktail, 40 µM PMSF and lysozyme). The cells were lysed and the cell debris was removed from the lysate by ultracentrifugation. Nedd4(HECT) was purified from the supernatant of the lysate as follows: the supernatant was loaded onto a $Ni^{2+}$-charged immobilized metal affinity column (equilibrated with 50 mM Tris, 100 mM NaCl, 20 mM imidazole, pH 7.5). Nedd4(HECT) was eluted via gradient elution from 20 to 250 mM imidazole. Nedd4(HECT)-containing fractions were pooled dialyzed overnight against 4 L of dialysis buffer (50 mM Tris, 100mM NaCl, pH 7.5, 1 mM TCEP), and $His_6$-TEV protease (1 mg) was added into the dialysis tubing with Nedd4(HECT) for proteolytic cleavage of the hexahistidine tag. After overnight dialysis, Nedd4(HECT) was subsequently purified through an additional round of $Ni^{2+}$-charged immobilized metal affinity (with buffers as stated above), and untagged Nedd4(HECT) was collected in the flow-through or from the wash while $His_6$-TEV protease was separated from the desired protein. Nedd4(HECT) was subsequently concentrated and aliquoted for storage at 4 °C, –20 °C (in 40% glycerol), and –80 °C.

<u>Nedd4 (full-length):</u> Nedd4 was purified according to our previously optimized method.[1] pGEX-Nedd4 (after two rounds of site-directed mutagenesis for insertion of hexahistidine affinity tag and TEV protease cleavage sequence) was transformed into BL21(DE3)-CodonPlus-RIL competent *E. coli.* Successful transformants were selected for by growth on ampicillin/chloramphenicol double selection plates. For expression, a starter culture (100 mL LB media) was inoculated from a streak of colonies on the selection plate and allowed to reach saturation by overnight growth at 37 °C. Expression cultures were inoculated from saturated starter culture (10 mL starter culture per 1 L LB media) and grown at 37 °C until $OD_{600}$ = 0.6 at which time expression was inducted with the addition of IPTG to a final concentration of 100 µM and grown overnight at 18 °C. After growth, cells were harvested and used immediately for purification. The cell pellet was resuspended in lysis buffer (50 mM Tris, 250 mM NaCl, pH 7.4 with 1X protease inhibitor cocktail, 40 µM PMSF and lysozyme). The cells were lysed and the cell debris was removed from the lysate by ultracentrifugation. $His_6$-GST-Nedd4 was purified from the supernatant of the lysate as follows: the supernatant was loaded onto glutathione agarose column (Genesee Scientific) that had been pre-equilibrated with 50 mM Tris, 250 mM NaCl, pH 7.4. The column was washed with 10 column volumes of buffer and $His_6$-GST-Nedd4 was eluted with 7 column volumes of elution buffer (50 mM Tris, 250 mM NaCl, pH 7.4 with 20 mM glutathione reduced). $His_6$-GST-Nedd4 containing fractions were pooled dialyzed overnight against 4 L of dialysis buffer (50 mM Tris, 100mM NaCl, pH 7.5, 1 mM β-mercaptoethanol), and $His_6$-TEV protease (1 mg) was added into the dialysis tubing with Nedd4 for proteolytic cleavage of the $His_6$-GST tag. After overnight dialysis, Nedd4 was subsequently purified through $Ni^{2+}$-charged immobilized metal affinity chromatography wherein the resin was pre-equilibrated with wash buffer (50 mM Tris, 250 mM NaCl, pH 7.4 with 20 mM imidazole). Dialysate was loaded onto the equilibrated column and untagged Nedd4 was collected in the flow-through and early in a 10 column volume wash. $His_6$-TEV and cleaved $His_6$-GST were eluted from the column over a gradient elution from 20 to 250 mM imidazole. Untagged full-length Nedd4 was subsequently concentrated and aliquoted for storage at 4 °C, –20 °C (in 40% glycerol).

<u>Ubiquitin:</u> Ubiquitin purification was based on a method previously described.[2] pET15-ubiquitin was transformed into BL21(DE3) competent *E. coli* and successful transformants were selected for by growth on ampicillin selection plates. For expression, a starter culture (100 mL 2xYT media) was inoculated from a streak of colonies on the selection plate and were grown at 37 °C until $OD_{600}$ = 0.6. Expression cultures were inoculated from starter culture (10 mL starter culture per 1 L 2xYT media) and grown at 37 °C until $OD_{600}$ = 0.6 at which time expression was inducted with the addition of IPTG to a final concentration of 400 µM and for four additional hours at 37 °C. After growth, cells were harvested and pellets were stored at –20 °C until use for purification. For purification, cell pellets were thawed on ice and resuspended in lysis buffer (50 mM Tris pH 7.6, 1 mM phenylmethylsulfonyl fluoride, 1X protease inhibitor cocktail (Bimake) and 0.4 mg/mL lysozyme). Cells were subsequently lysed and cell debris was removed by centrifugation (Sorvall SS-34, 8000 rpm, 20 min at 4 °C). Supernatant was transferred to a cooled beaker on ice and perchloric acid (70%, 0.35 mL per 50 mL lysis buffer used) was added while stirring. Solution was centrifuged (Sorvall SS-34, 8000 rpm, 20 min at 4 °C) and supernatant was dialyzed against 2L dialysis buffer (50 mM ammonium acetate, pH 4.5) overnight. Dialysis tubing was transferred to 2L fresh dialysis buffer and dialyzed for an additional few hours. After dialysis, ubiquitin was further purified by ion exchange chromatography using SP Sepharose (column equilibrated with 50 mM ammonium acetate, pH 4.5) and was eluted over a gradient elution from 0 to 0.5 M NaCl in 50 mM ammonium acetate. Fractions were analyzed by SDS-PAGE and fractions containing ubiquitin were pooled and concentrated. Samples were lyophilized or buffer exchanged into desired assay reaction buffers.

*Site-directed mutagenesis*

All site-directed mutagenesis was performed following Q5 site-directed mutagenesis kit and published manufacturer's protocol (New England Biolabs). Two rounds of mutagenesis were performed for insertion of a hexahisitidne sequence and a cleavage sequence for TEV protease into the pGEX-

Nedd4 plasmid. Primers used for mutagenesis were designed with NEBaseChanger (New England Biolabs) and are described in Supplemental Table 2. Mutagenesis was confirmed by Sanger sequencing as performed by Eton Biosciences.

*Small molecule synthesis*

Materials: Unless otherwise stated, all reagents were purchased from either Sigma-Aldrich or Oakwood Chemicals and were used as received. $CDCl_3$ was purchased from Cambridge Isotope Laboratories and used as received.

Characterization: NMR spectra were obtained using Bruker Advance Neo - 500 MHz multinuclear spectrometer (Duke Chemistry NMR Facility) operating at 500, MHz for $^1H$ NMR and 126 MHz for $^{13}C$ NMR and are reported as chemical shifts (δ) in parts per million (ppm). Spectra were referenced internally according to residual solvent signals ($^1H$, $CDCl_3$ 7.26 ppm; $^{13}C$, $CDCl_3$ 77.0 ppm). Data for NMR spectra use the following abbreviations to describe multiplicity: s, singlet; br s, broad singlet; d, doublet; t, triplet; dd, doublet of doublets; td, triplet of doublets; tt, triplet of triplets; ddd, doublet of doublets of doublets; dddd, doublet of doublets of doublets of doublets; ddt, doublet of doublets of triplets; app dd, apparent doublet of doublets; m, multiplet. Coupling constants (J) are reported in units of hertz (Hz). High-resolution mass spectra (HRMS, m/z) were recorded on an Agilent 6224 LC/MS-TOF spectrometer using electrospray ionization (ESI, Duke University Department of Chemistry Shared Instrumentation Facility).

Preparation of *N*-arylbenzimidaozle (NAB) derivatives: *N*-arylbenzimidazole derivatives NAB2 and NAB17 were synthesized according to previously reported procedures[3] as described below (Supplemental Scheme 1). Characterization via NMR and HRMS are consistent with previously reported characterizations.

*Supplemental Scheme 1: Synthesis of N-(2-chlorobenzyl)-4-fluoro-3-nitrobenzamide*

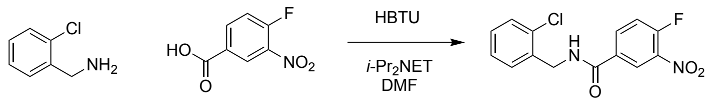


**N-(2-chlorobenzyl)-4-fluoro-3-nitrobenzamide.** 2-Chlorobenzylamine (0.71 mL, 6.0 mmol) was added slowly to a stirred solution of 4-fluoro-3-nitrobenzoic acid (925 mg, 5.0 mmol), HBTU (1.90 g, 5.0 mmol), *N,N*-diisopropylethylamine (1.05 mL, 6.0 mmol), and DMF (10 mL) at room temperature. After 30 min, the solution was diluted with ethyl acetate (50 mL) and washed sequentially with water, 1M HCl (aq) (3X), 1M KOH (aq) (3X), and brine. The organic layer was dried over sodium sulfate, filtered, and concentrated under reduced pressure using a rotary evaporator. Purification of the residue by silica gel chromatography (10-45% EtOAC/Hexanes; material loaded using toluene) provided the title compound as a yellow solid (0.77 g, 50% yield). $^1H$ NMR (500 MHz, $CDCl_3$) δ 8.46 (dd, $J$ = 7.0, 2.3 Hz, 1H), 8.11 (ddd, $J$ = 8.8, 4.1, 2.3 Hz, 1H), 7.50 – 7.34 (m, 4H), 7.28 (m, 2H), 6.60 (s, 1H), 4.75 (d, $J$ = 5.9 Hz, 2H). $^{13}C$ NMR (126 MHz, $CDCl_3$) δ 164.30, 157.24 (d, J = 270.9), 134.96, 134.67 (d, J = 8.8), 133.73, 131.18 (d, J = 3.8) 130.28, 129.72, 129.33, 127.25, 125.2, 118.95 (d, J = 21.4), 42.43. HRMS (ESI-TOF) *m/z* calcd for $C_{14}H_{10}ClFN_2O_3$ $[M+H]^+$ 309.0437, found 309.0445.

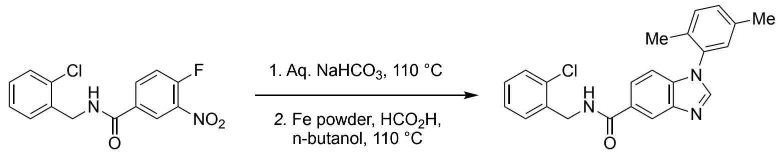


**N-(2-chlorobenzyl)-1-(2,5-dimethylphenyl)-1H-benzo[d]imidazole-5-carboxamide (NAB2).** A screw-top test tube containing N-(2- chlorobenzyl)-4-fluoro-3-nitrobenzamide (0.96 mmol, 300 mg), sodium bicarbonate (1.92 mmol, 161 mg), 2,5-dimethylaniline (1.92 mmol, 240 $\mu$L), and water (2 mL/mmol benzamide) was sealed with a Teflon screw cap, placed in a preheated oil bath at 110 °C, and stirred for 24 h. The reaction was cooled to room temperature and the mixture was poured onto ethyl acetate (sufficient volume for all solid to be dissolved). The solution washed sequentially with 1M HCl (aq) (3X), water (3X), and brine (3X). The organic layer was dried over sodium sulfate, filtered, and concentrated using reduced pressure via a rotary evaporator. The crude nitroaniline was subsequently dissolved in n-butanol (1.0 mL/mmol benzamide, 0.96 mL), transferred to a second screw-top test tube, and formic acid (1.0 mL/mmol benzamide, 0.96 mL), iron powder (9.6 mmol, 536 mg), and concentrated HCl (0.1 mL/mmol benzamide, 96 $\mu$L) were added to the solution. The test tube was sealed with a Teflon screw cap and stirred for 1 h at 110 °C. After cooling to room temperature, the mixture was poured onto a mixture of ethyl acetate and saturated $NaHCO_3$ (1:6 ratio) in a separatory funnel. The mixture was shaken with venting and solid sodium bicarbonate added until pH~12. The layers were separated and the organic layer was washed with water (3X), dried and concentrated under reduced pressure. The residue was purified by silica gel chromatography using 40-100% EtOAC/Hexanes (material loaded using chloroform) to afford NAB2 as a white solid (194.1mg, 52% yield). $^1H$ NMR (500 MHz, $CDCl_3$) δ 8.27 (s, 1H), 8.01 (s, 1H), 7.83 (dd, J = 8.5, 1.7 Hz, 1H), 7.52 (dd, J = 7.0, 2.4 Hz, 1H), 7.40 (dd, J = 7.0, 2.2 Hz, 1H), 7.31 (d, J = 7.9 Hz, 1H), 7.29 – 7.22 (m, 3H), 7.18 (d, J = 8.5 Hz, 1H), 7.11 (s, 1H), 6.71 (bs, 1H), 4.79 (d, J = 5.9 Hz, 2H), 2.40 (s, 3H), 2.03 (s, 3H). $^{13}C$ NMR (126 MHz, $CDCl_3$) δ 167.85, 144.65, 143.01, 137.45, 137.01, 136.00, 134.14, 133.78, 132.04, 131.53, 130.55, 130.37, 129.67, 129.27, 129.03, 128.10, 127.27, 123.49, 119.26, 110.91, 42.24, 20.89, 17.19. HRMS (ESI-TOF) *m/z* calcd for $C_{23}H_{20}ClN_3O$ $[M+H]^+$ 390.1368, found 390.1371.

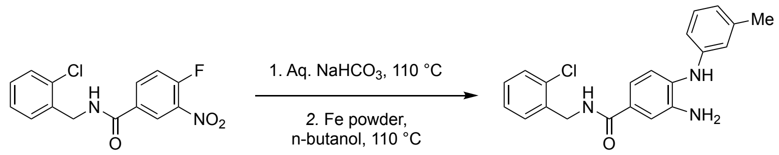


**3-amino-N-(2-chlorobenzyl)-4-(m-tolylamino)benzamide (NAB17).** A screw-top test tube containing N-(2- chlorobenzyl)-4-fluoro-3-nitrobenzamide (0.32 mmol, 100 mg), sodium bicarbonate (0.64 mmol, 53 mg), 3-methylaniline (0.64 mmol, 68 $\mu$L), and water (2 mL/mmol benzamide) was sealed with a Teflon screw cap, placed in a preheated oil bath at 110 °C, and stirred for 24 h. The reaction was cooled to room temperature and the mixture was poured onto ethyl acetate (sufficient volume for all solid to be dissolved). The solution washed sequentially with 1M HCl (aq) (3X), water (3X), and brine (3X). The organic layer was dried over sodium sulfate, filtered, and concentrated using reduced pressure via a rotary evaporator. The crude nitroaniline was subsequently dissolved in n-butanol (1.0 mL/mmol benzamide, 0.32 mL), transferred to a second screw-top test tube, iron powder (3.2 mmol, 179 mg), and concentrated HCl (0.1 mL/mmol benzamide, 0.032 mL) were added to the solution. *NOTE: formic acid was omitted from this reaction mixture to prevent closure of the heterocycle.* The test tube was sealed with a Teflon screw cap and stirred for 1 h at 110 °C. After cooling to room temperature, the mixture was poured onto a mixture of ethyl acetate and saturated $NaHCO_3$ (1:6 ratio) in a separatory funnel. The mixture was shaken with venting and solid sodium bicarbonate added until pH~12. The layers were separated and the organic layer was washed with water (3X), dried and concentrated under reduced pressure. The crude product was worked up as described and purified by silica gel chromatography using 40-100% EtOAC/Hexanes (material loaded using chloroform) to afford NAB17 as an off-white solid (72.7 mg, 62.1%). $^{1}H$ NMR (500 MHz, $CDCl_3$) δ 7.50 – 7.44 (m, 1H), 7.39 (dd, *J* = 7.2, 2.2 Hz, 1H), 7.31 (d, *J* = 2.0 Hz, 1H), 7.25 – 7.20 (m, 2H), 7.17 – 7.08 (m, 3H), 6.74 (d, *J* = 7.6 Hz, 1H), 6.69 (d, *J* = 6.2 Hz, 2H), 6.49 (m, 1H), 5.33 (s, 1H), 4.72 (d, *J* = 6.0 Hz, 2H), 2.29 (s, 3H), 2.01 (d, *J* = 1.2 Hz, 1H). $^{13}C$ NMR (126 MHz, $CDCl_3$) δ : 167.30, 143.44, 139.53, 139.46, 135.97, 133.86, 133.81, 130.58, 129.70, 129.56, 129.42, 129.12, 127.32, 121.90, 120.57, 118.06, 117.93, 115.86, 114.44, 42.13, 21.66. HRMS (ESI-TOF) *m/z* calcd for $C_{21}H_{20}ClN_3O$ $[M+H]^+$ 366.1368, found 366.1376.

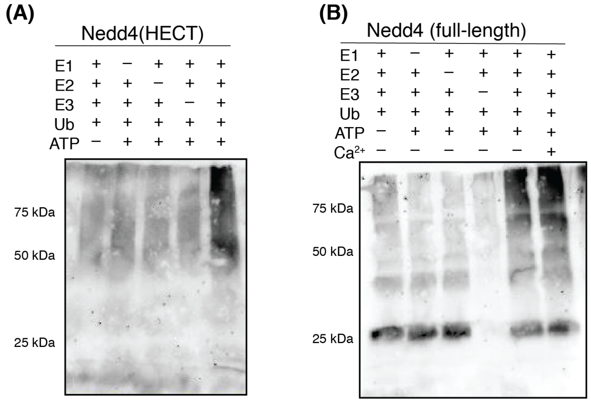


**Supplemental Figure 1. Validation of ligase activity by *in vitro* ubiquitination assay** Immunoblotting-based activity assay (IB: anti-ubiquitin, 1:2000) of Nedd4 ligase activity shows that ubiquitin chains (free chains, E3 autoubiquitination, or substrate ubiquitination) are only formed in the presence of catalytically active E1 activating enzyme, E2 conjugating enzyme and E3 ligase. Assay was performed for 60 minutes at 37°C with E1 (100 nM), E2 (1 $\mu$M), E3 (5 $\mu$M), ubiquitin (100 $\mu$M) and ATP (2 mM). Both **(A)** Nedd4(HECT) and **(B)** full-length Nedd4, which were accessed as recombinant enzymes, are active.

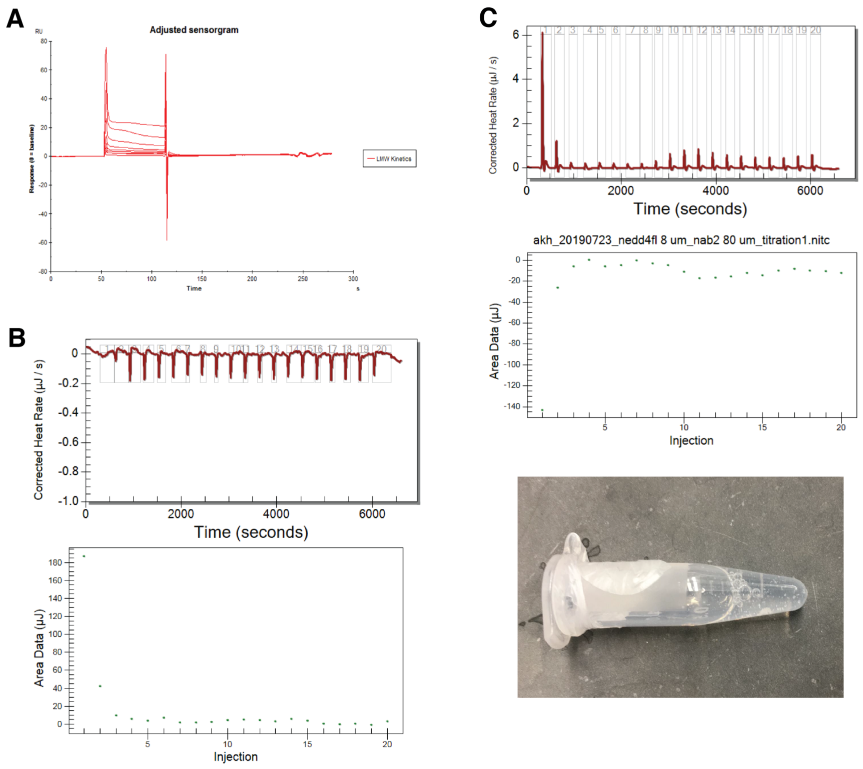


**Supplemental Figure 2. SPR and ITC experiments to characterize NAB2 binding to full-length Nedd4 and Nedd4(HECT)** Investigation of NAB2 binding to Nedd4(HECT) by **(A)** surface plasmon resonance did not produce a canonical binding curve when Nedd4(HECT) was immobilized on the SPR surface and exposed to a gradient of NAB2 from 0 to 100 $\mu$M. **(B)** Further analysis of NAB2 binding to Nedd4(HECT) was conducted via isothermal titration calorimetry (ITC) in which NAB2 (80 $\mu$M) was titrated into Nedd4(HECT) (8 $\mu$) over 20 injections over 100 minutes. **(C)** ITC analysis of NAB2 binding to full-length Nedd4 resulted in some heat release but did not show a canonical ITC binding isotherm for a 1:1 ligand:enzyme interaction. Additionally, a large degree of precipitate was present in the protein sample when the sample was removed after the titration experiment (bottom), indicating that the effective protein concentration decreased during the course of the experiment. Precipitation persisted even with the addition of stabilizing agents like glycerol to the ITC sample preparation buffer. Due to this experimental artifact, alternative thermodynamic measures of ligand binding were pursued.

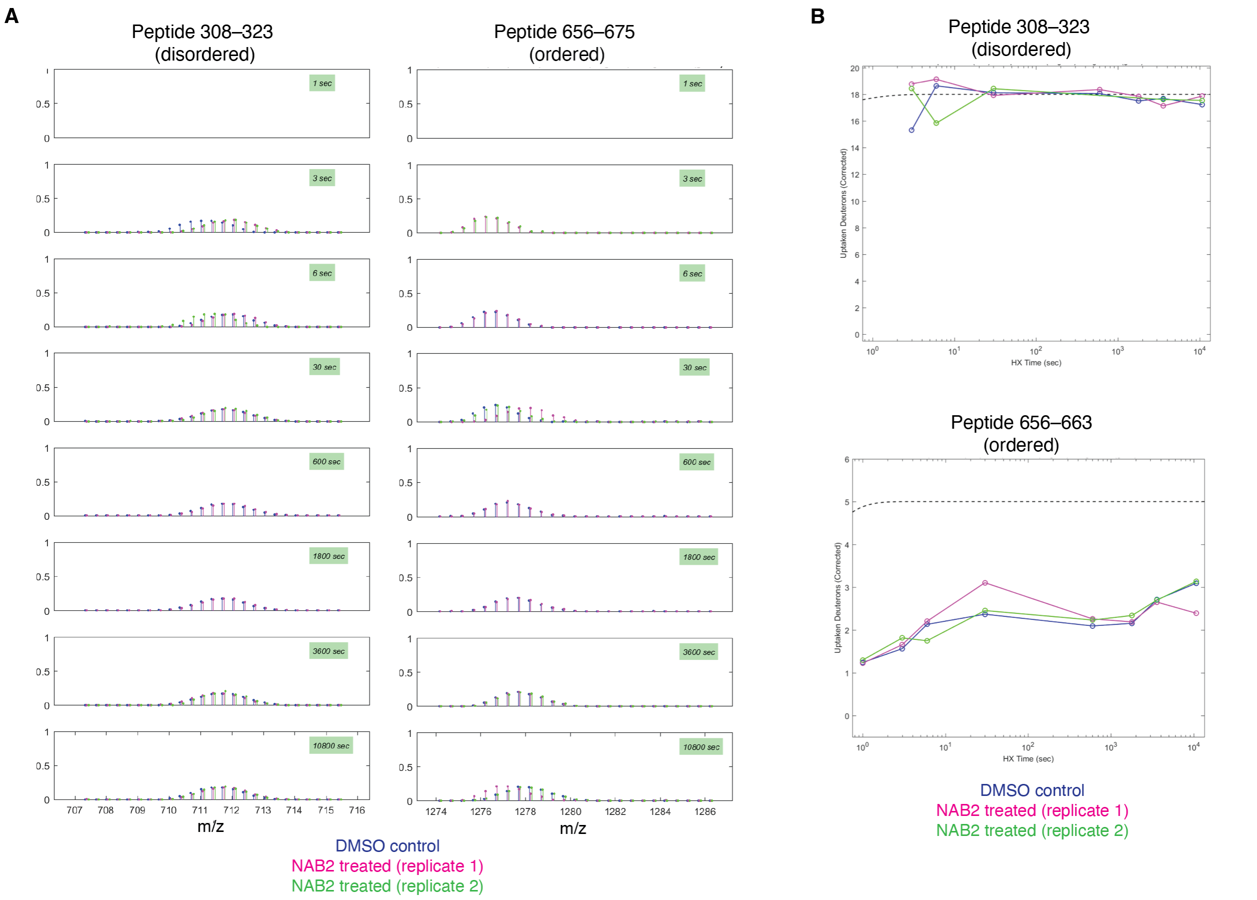


**Supplemental Figure 3. Peptide-level detection of deuterium uptake over time in hydrogen-deuterium exchange mass spectrometry (HDX-MS) analysis of Nedd4** HDX-MS analysis was conducted with ExMS2 software wherein deuterium uptake at each timepoint is visualized at a peptide level. As the rate of deuterium uptake is dependent upon the relative order of the protein, structured and unstructured regions of Nedd4 display very different deuterium uptake profiles. **(A)** Centroid and **(B)** time-course plots from HDX-MS analysis for peptides from ordered and disordered of Nedd4 are shown. Represented peptides are located in a linker region between WW domains 1 and 2 (peptide 30–323) or in the structured HECT domain (peptides 656–675 and 656–663). As demonstrated by the representative images, peptides in unstructured regions may be fully deuterated (as determined by comparison to “all deuterium” control) even at the shortest time regimes while peptides in ordered regions may not reach full deuteration.

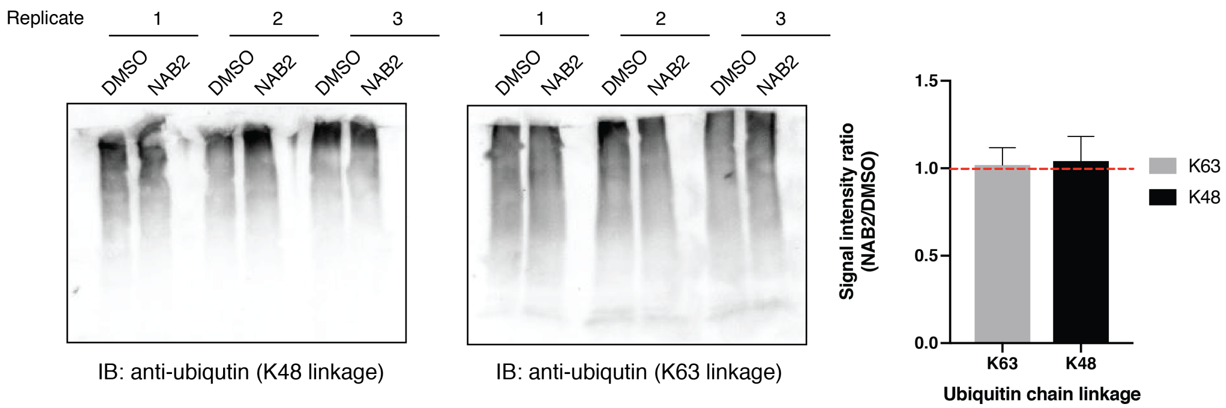


**Supplemental Figure 4. NAB2 treatment does not alter Nedd4 ubiquitination linkage specificity** Immunoblotting assays indicate that NAB2 treatment of Nedd4 (full-length, recombinant) does not induce a change in the ubiquitination linkage specificity of the ligase. Assays were conducted for 60 min at 37 °C in the presence of E1 (100 nM), E2 (1 $\mu$M), E3 (2 $\mu$M), and ubiquitin (100 $\mu$M) in the presence of ATP (2 mM) and either NAB2 (20 $\mu$M) or DMSO control. Ubiquitin chain linkage was detected by immunoblotting with primary antibodies specific to either K48 or K63 linkages (Abcam, 1:2000 dilution) as K48 and K63 linkages are the predominant chain types produced by Nedd4. Immunoblotting signal was quantified as the ratio of signal intensity from NAB2 treated samples vs DMSO control wherein a ratio of one indicates no change in linkage specificity.

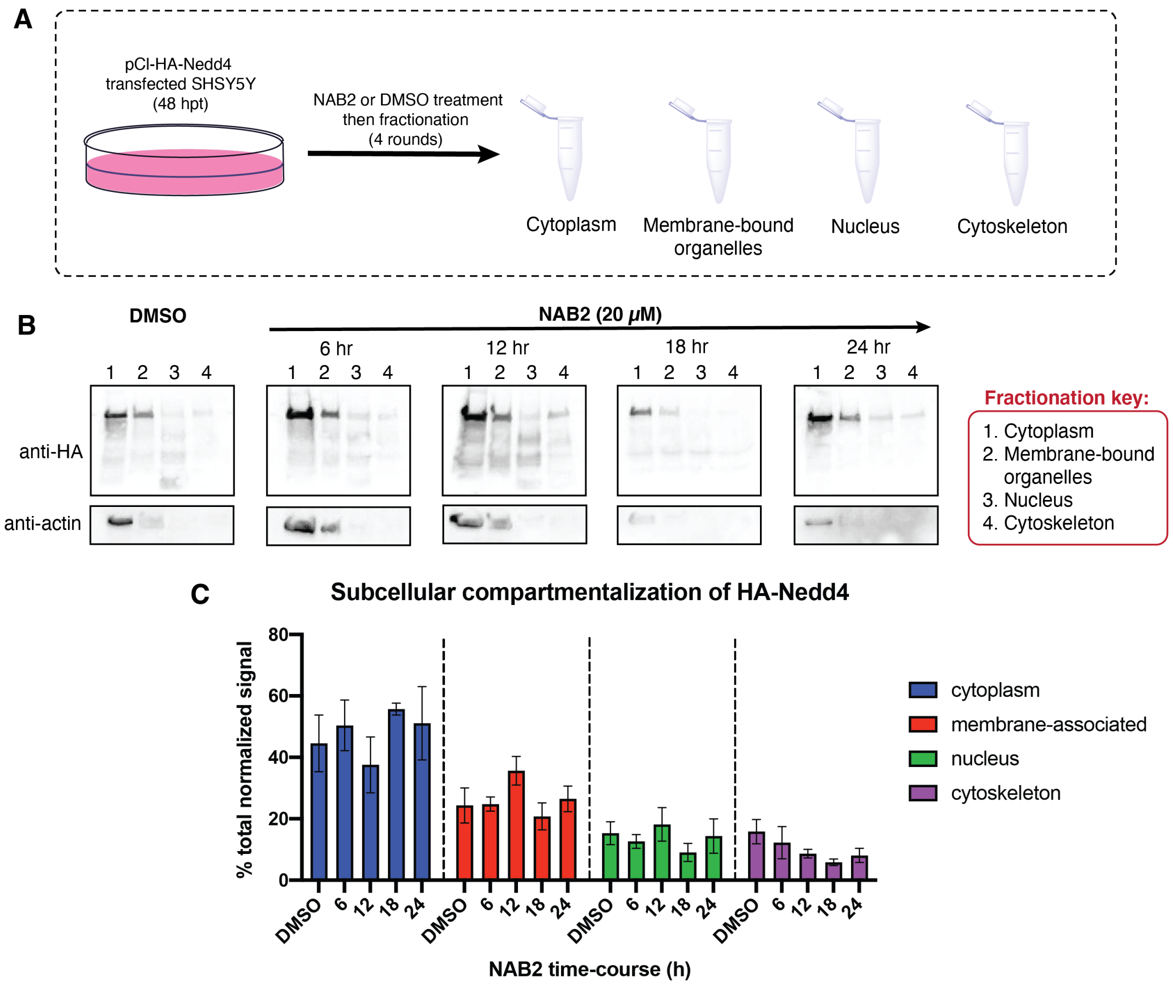


**Supplemental Figure 5. Immunoblotting for detection of Nedd4 subcellular compartmentalization** Subcellular fractionation shows that Nedd4 compartmentalization is not altered upon treatment of NAB2 relative to a DMSO control. **(A)** Experimental workflow (top) was as follows: wild-type SHSY5Y cells were transfected with pCI-HA-Nedd4 and treated with DMSO or NAB2 at 48 hpt. Following treatment, cells were incubated for the indicated time (6, 12, 18 or 24h). Cells were trypsinized and were subsequently fractionated with the ProteoExtract subcellular fractionation kit (Millipore Sigma) according to the protocol for suspension cells. Samples were separated by SDS-PAGE and were transferred to activated PVDF membrane for analysis by immunoblotting with anti-HA and anti-actin followed by HRP-conjugated secondary antibodies. **(B)** Representative immunoblots for detection of NAB2-dependent changes in Nedd4 subcellular compartmentalization are shown (bottom). **(C)** Quantification of fractionation experiments indicate that NAB2 treatment does not significantly alter Nedd4 compartmentalization over time. The fractionation experiment was completed in triplicate and replicate data is shown as the mean + s.e.m. of the replicates.

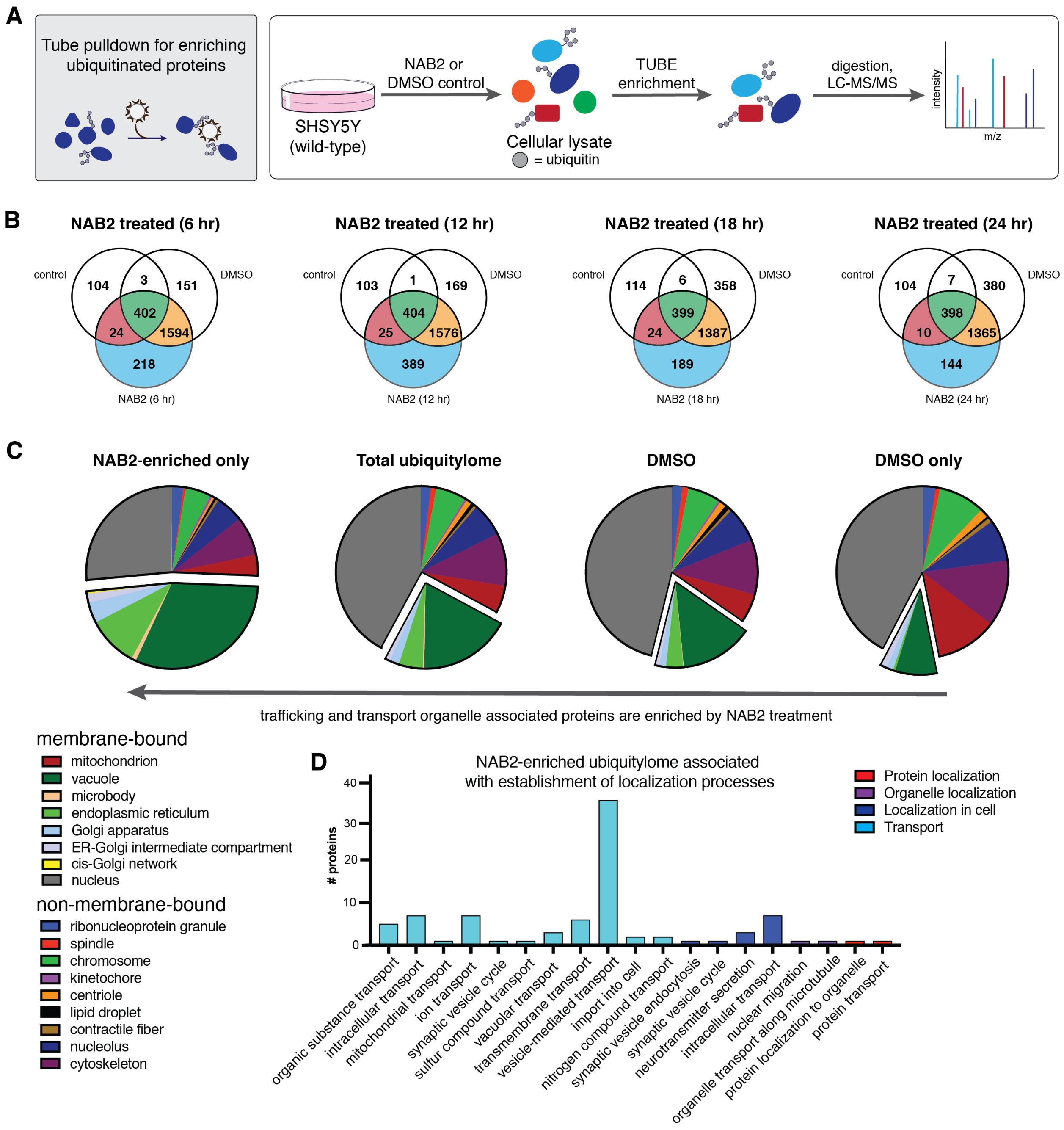


**Supplemental Figure 6. Qualitative proteomic analysis of the enriched ubiquitylome indicates that NAB2 treatment increases the portion of trafficking and transport proteins in the ubiquitylome. (B)** Samples were prepped according to the workflow show in Figure 4A and were subsequently subjected to analysis by LC-MS/MS. Qualitative analysis of the ubiquitylome enriched from neuroblastoma-derived SHSY5Y cells shows that NAB2 treatment produces a unique pool of ubiquitinated proteins relative to DMSO control (blue). Across all samples, there were 2841 proteins identified by qualitative proteomics. Of that set of proteins, there are 532 unique, non-redundant proteins that are present only after NAB2 treatment. **(C)** Functional annotation of identified proteins with the PANTHER functional classification algorithm reveal that NAB2 treatment increases the portion of proteins associated with trafficking and transport organelles in the ubiquitylome relative to DMSO control. **(D)** Further annotation of proteins enriched by NAB2 treatment indicates that a majority of NAB2-dependent hits are involved in specific vesicle-mediated transport processes.

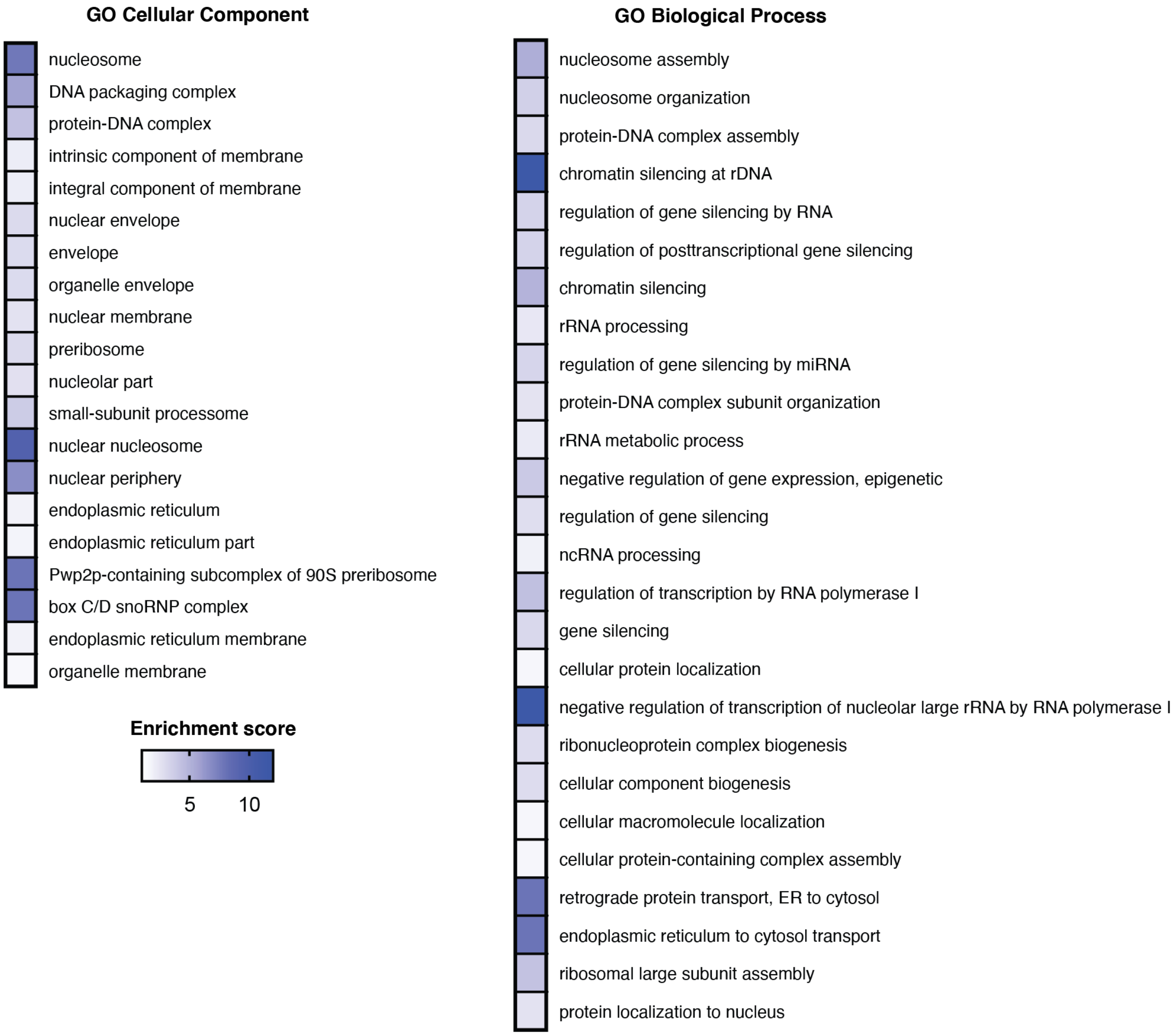


**Supplemental Figure 7. Bioinformatic analysis of α-synuclein-dependent ubiquitylome via Gene Ontology** (GO) enrichment Proteins in the enriched ubiquitylome that were significantly altered by α-synuclein toxicity (α-synuclein-A53T OE SHSY5Y) relative to control cells (WT SHSY5Y) were analyzed using the GOrilla[4] program for Gene Ontology enrichment analysis (http://cbl-gorilla.cs.technion.ac.il/). GO enrichment analysis shows that α-synuclein toxicity induces ubiquitination of proteins in various cellular compartments (left) and biological processes (right) relative to the total ubiquitylome. Enrichment score (calculated with GOrilla[4]) was calculated by comparison of an unranked target list (significant proteins in α-synuclein-A53T vs WT) compared to a background list of genes (all proteins identified in the ubiquitylome) using the default parameters.

**Supplemental Table 1.** Plasmids used for recombinant protein expression or mammalian cell transfection, acquired via Addgene plasmid repository

| Plasmid | GenBankID | Expression System | Addgene Plasmid # | Depositor | Modified? |
|---|---|---|---|---|---|
| pET15-UbE2D1 (Ubch5a) | NM_003338 | Bacterial | 15782 | Wade Harper | No |
| pGEX-Nedd4 | NM_006154.2 | Bacterial | 45043 | Shiaw-Yih Lin | Yes |
| pET15-ubiquitinWT | AAA72608.1 | Bacterial | 12647 | Rachel Klevit | No |
| pET28-MHL-5c7J (Nedd4(HECT)) | NP_006154.2 (residues 520-900) | Bacterial | 69965 | Cheryl Arrowsmith | No |
| pCl-HA-Nedd4 | NM_006154.2 | Mammalian | 27002 | Joan Massague | No |

**Supplemental Table 2.** Primers used for Q5 site-directed mutagenesis

| Plasmid | Q5 modification | Desired sequence | Primer | | Location of modification |
|---|---|---|---|---|---|
| pGEX-Nedd4 | Insertion | ENLYFQG | F | 5'-CACCACCCACTCCCCTATACTAGGTATTG-3' | Between residues 1 and 2 of ORF |
| | | | R | 5'-ATGATGATGCATGAATACTGTTTCCTGTG-3' | |
| pGEX-Nedd4 | Insertion | HHHHHH | F | 5'-TTTTCAGGGCGCCCTTATGGCAACTTGC-3' | Between residues N230 and S231 of ORF |
| | | | R | 5'-TACAGGTTTTCGAATTCGGGGATCCCACG-3' | |

**Supplemental Table 3.** List of gene names of previously identified Nedd4 interactors compiled from various protein-protein interaction databases. Bold indicates proteins which are known Nedd4 interactors that were identified in the TUBE-coupled quantitative proteomics dataset, and red proteins are significantly altered in the proteomics dataset upon induction of α-synuclein toxicity relative to wild-type cells.

| | | | | | | |
|---|---|---|---|---|---|---|
| 43717 | CUEDC1 | **HGS** | MGRN1 | PTGFR | SLC38A3 | TPCN2 |
| A0A024R8L2 | **CUL1** | HMCES | MLANA | PTK2B | SLC5A7 | **TRAF6** |
| **ABCB1** | CUL5 | **HNRNPK** | **MLH1** | **RAC1** | SLC6A3 | TRIM44 |
| ABCG1 | DALLY | **HNRNPL** | MOB3A | **RAD51AP1** | SLC9A1 | TRIM52 |
| ABL1 | DCUN1D1 | **HNRNPM** | MRAP2 | **RAF1** | SMAD1 | TRPV6 |
| ABL2 | DDIT4 | **HNRNPU** | MRPL19 | RALBP1 | **SMAD2** | TSC1 |
| ADRB2 | **DDX3X** | **HNRNPUL1** | MST4 | **RANBP10** | SMAD3 | **TSG101** |
| AGAP2 | **DDX54** | HSF1 | MTMR4 | RAP2A | **SMAD5** | TSPAN3 |
| AKT1 | **DHX30** | IFITM3 | MYCN | RAPGEF2 | SMAD6 | TSTA3 |
| AKT3 | **DIAPH1** | IFNGR1 | MYZAP | RAPGEF6 | SMAD7 | TTYH2 |
| AMOT | DMD | IGF1R | N4BP1 | RASGEF1A | **SMARCC1** | TTYH3 |
| AMOTL1 | DRD2 | IL2RB | N4BP3 | RASL11B | SMO | **UBAP2L** |
| AMPD2 | DVL1 | IL2RG | NDFIP1 | **RBCK1** | SMOI | **UBB** |
| AMPH | DYRK4 | ILDR1 | NDFIP2 | RBX1 | SMURF2 | UBC |
| ANKRD13D | EBAG9 | **IMPDH2** | **NEDD4** | **RER1** | **SNCA** | UBC8 |
| ANXA13 | EGFR | INO80 | NEDD4L | RET | SP140L | UBE2D1 |
| ANXA9 | ENTPD7 | IRS1 | NFATC2 | RFT1 | SPANXN3 | **UBE2D2** |
| AP1G2 | EPHA5 | IRS2 | NFE2 | RIPK1 | SPRY1 | **UBE2D3** |
| AQP2 | EPRS | ISG15 | **NHP2** | RNF11 | SPRY2 | UBE2E1 |
| **ARID1A** | **EPS15** | **ITCH** | **NSRP1** | RNF7 | SPT23 | UBE2L3 |
| ARMH4 | ERBB3 | IZUMO1 | NTRK1 | RPAP2 | **SQSTM1** | **UBE2M** |
| ARRB1 | **ERBB4** | **JUN** | **NUDT21** | **RPAP3** | SRC | UBI4 |
| **ARRB2** | ERG | KCNAB1 | **NUMB** | **RPL18A** | SRMS | UBOX5 |
| ARRDC1 | ERMN | KCNAB2 | OCLN | RPO21 | **SRSF7** | UL56 |
| ARRDC3 | ERRFI1 | KCNJ16 | OPN4 | **RPS27A** | STK24 | ULK1 |
| ARRDC4 | FAM189A2 | KCNQ1 | OSTCL | **RPS3A** | **STK25** | UQCRH |
| **ASPSCR1** | FAM189B | KDM2A | P0CK58 | **RPS6KA3** | STK26 | URI1 |
| AURKC | FAM8A1 | KDM6B | PARP16 | RPS6KA4 | STK31 | US9 |
| **BAIAP2** | **FBXL15** | KIAA1598 | PAX7 | RPS6KB1 | STK4 | **USP10** |
| BDKRB1 | **FBXL18** | KIF26B | **PDCD6IP** | **RPSA** | SULF1 | USP2 |
| BEAN1 | FES | KIFC3 | PDGFRB | **RTF1** | SUSD6 | **USP36** |
| BECN1 | FGF12 | KUZ | **PDLIM7** | RUNX1 | SUV420H2 | VIF |
| BFRF1 | FGF21 | LAMP2 | PI4K2A | **RUVBL1** | SYK | WBP1 |
| BMPR1A | FGFR1 | LAPTM4A | **PIP5K1A** | SAAL1 | SYT1 | **WBP2** |
| BRCA2 | FGFR2 | LAPTM4B | PIP5K1C | SALS | TAF1B | WEE1 |
| BT.104932 | FKBP3 | LAPTM5 | **PKM** | SAMSN1 | **TAX1BP1** | WIBG |
| BTRC | FLT1 | LATS1 | **PKN2** | SAV1 | TBC1D7 | **XPO1** |
| C11ORF63 | FLT4 | LATS2 | **PLK1** | **SCAMP3** | **TBK1** | YES1 |
| C16ORF58 | **FMR1** | LDLRAD4 | PLK2 | SCN5A | TCEANC | **YOD1** |
| C3ORF37 | **FXR1** | LITAF | PMEPA1 | SCN8A | TCP11L1 | **YWHAB** |
| **CAD** | FYN | LMP2 | **POLR1C** | SCNN1A | TEAD2 | **YWHAE** |
| CALCOCO1 | GABARAP | LRRIQ1 | **POLR2A** | SCNN1B | TEN-M | **YWHAH** |
| **CAMK1D** | GABARAPL1 | LRRK1 | **POLR2B** | SCNN1G | TGFBR2 | **YY1** |
| CAMK4 | GABARAPL2 | **LUC7L2** | **POLR2C** | SDHA | **THOC1** | ZDHHC7 |
| CAMKK2 | GAG | MANSC1 | **POLR2E** | SERTAD1 | **THRAP3** | ZFP36L2 |
| CBLB | GBA | MAP1LC3A | **POLR2M** | SFTPC | TMEM171 | |
| **CCNH** | GGA3 | MAP1LC3B | **POLR3A** | SGK1 | TMEM231 | |
| CCR2 | GINM1 | MAP1LC3C | PPARG | SGK2 | TMEM255A | |
| **CD2BP2** | GJA1 | MAP2 | PRKG2 | SHANK3 | TMEM51 | |
| CD83 | GLIS3 | MAP3K2 | PRKX | SHISA6 | TMEM52B | |
| CDC25C | GRB10 | MAP3K3 | PROM1 | SIVA1 | TMEM92 | |
| **CDK5** | GRIA1 | MAP3K5 | **PRPF8** | SKIL | TNFAIP3 | |
| CDK5R1 | GRIN2A | MAP4K5 | PRR16 | **SKP1** | **TNFRSF10B** | |
| CFTR | GRIN2D | MAPK14 | PRRG1 | SLC11A2 | TNFRSF19 | |
| CLIC2 | GRK4 | MAPKAPK3 | PRRG2 | SLC12A3 | **TNIK** | |
| **CLK3** | GRK7 | **MARK2** | PRRG3 | SLC1A1 | TNK2 | |
| COMM | GUCD1 | **MARK4** | **PSMD4** | SLC20A1 | **TOM1** | |
| **CPSF1** | H1F0 | MB21D1 | PTC | SLC22A6 | **TOM1L2** | |
| **CPSF6** | H3F3A | MDM2 | PTCH1 | SLC22A8 | **TP53BP2** | |
| **CSNK1D** | HEPACAM2 | MED28 | PTEN | SLC23A2 | TP73 | |
